# Inhibitory temporo-parietal effective connectivity is associated with explicit memory performance in older adults

**DOI:** 10.1101/2022.12.22.521668

**Authors:** Björn H. Schott, Joram Soch, Jasmin M. Kizilirmak, Hartmut Schütze, Anne Assmann, Anne Maass, Gabriel Ziegler, Magdalena Sauvage, Anni Richter

**Affiliations:** German Center for Neurodegenerative Diseases (DZNE), Göttingen, Germany; Bernstein Center for Computational Neuroscience (BCCN), Berlin, Germany; Leibniz Institute for Neurobiology (LIN), Magdeburg, Germany; German Center for Neurodegenerative Diseases (DZNE), Magdeburg, Germany; Otto von Guericke University, Medical Faculty, Magdeburg, Germany; Center for Behavioral Brain Sciences (CBBS), Magdeburg, Germany; Department of Psychiatry and Psychotherapy, University Medical Center Göttingen, Göttingen, Germany; German Center for Mental Health (DZPG), Magdeburg, Germany; Center for Intervention and Research on adaptive and maladaptive brain Circuits underlying mental health (C-I-R-C) Jena-Magdeburg-Halle, Magdeburg, Germany

**Author notes:** Address for correspondence: PD Dr. Dr. Björn Hendrik Schott Leibniz Institute for Neurobiology Brenneckestr. 6, 39118 Magdeburg, Germany, /.

**Keywords:** subsequent memory effect, episodic memory, dynamic causal modeling, fMRI, cognitive aging, precuneus, hippocampus

## Abstract

Successful explicit memory encoding is associated with inferior temporal activations and medial parietal deactivations, which are attenuated in aging. Here we used Dynamic Causal Modeling (DCM) of functional magnetic resonance imaging data to elucidate effective connectivity patterns between hippocampus, parahippocampal place area (PPA) and precuneus during encoding of novel visual scenes. In 117 young adults, DCM revealed pronounced activating input from the PPA to the hippocampus and inhibitory connectivity from the PPA to the precuneus during novelty processing, with both being enhanced during successful encoding. This pattern could be replicated in two cohorts (N = 141 and 148) of young and older adults. In both cohorts, older adults selectively exhibited attenuated inhibitory PPA-precuneus connectivity, which correlated negatively with memory performance. Our results provide insight into network dynamics underlying explicit memory encoding and suggest that age-related differences in memory-related network activity are, at least partly, attributable to altered temporo-parietal neocortical connectivity.

## 1. Introduction

One of the core questions in the cognitive neuroscience of episodic memory is why some experiences are encoded into stable memory traces that can subsequently be retrieved, while other experiences are forgotten. In functional magnetic resonance imaging (fMRI) studies of episodic memory, this question is commonly investigated using the so-called subsequent memory approach ^1^. Since its first application to fMRI ^2,3^, numerous studies have convergingly shown that successful encoding robustly engages the medial temporal lobe (MTL) and particularly the hippocampus (HC), as well as inferior temporo-occipital regions like the parahippocampal place area (PPA), prefrontal, and lateral parietal cortices (for a meta-analysis, see ^4^). On the other hand, structures of the cortical midline like the precuneus (Prc) and the medial prefrontal cortex (mPFC), which form part of the Default Mode Network (DMN), are associated with relative deactivations during successful versus unsuccessful encoding ^4,5^.

Memory performance almost invariably declines with increasing age ^6^, and this decline is accompanied by characteristic alterations in memory-related network activations, including a reduced activation in inferior temporal cortices like the PPA and a reduced deactivation or even atypical activation of DMN, and particularly cortical midline structures such as the Prc ^5,7,8^). Despite the robustness of these findings, the neural mechanisms underlying the relatively higher DMN activity during encoding still remain unclear. They may reflect increased reliance on DMN-dependent cognitive processes like self-referential or prior knowledge-dependent information during encoding ^5^ or a reduced ability to suppress unwanted DMN activity, reflecting lower processing efficiency or specificity ^9–11^.

One potential mechanism mediating encoding-related hyperactivation of DMN structures in older adults could be increased excitatory or decreased inhibitory connectivity within the temporo-parietal memory network. The HC is highly interconnected with distributed neocortical regions ^12^, and previous studies have highlighted both the importance of hippocampal-neocortical connectivity for successful memory formation ^13–18^ and the susceptibility of memory-related network connectivity to age-related alterations ^13,19^.

Most functional connectivity measures, however, do not normally allow inferences about the directionality of information flow or about its excitatory versus inhibitory nature. Such information can be deduced from analyses of *effective* connectivity, such as Granger causality analysis (GCA) or Dynamic Causal Modeling (DCM) ^20^, but these approaches have thus far rarely been used in the context of memory encoding in older adults. Results from previous studies suggest that prefrontal-hippocampal effective connectivity becomes less task-specific in older adults with memory impairment ^21^ and that age-related deficits in HC-dependent navigation learning may be explained by higher HC excitability in older adults ^22^. However, no studies have thus far characterized the encoding-related information flow between the HC and the brain structures that show the most prominent age-related under-recruitment (i.e., inferior temporal structures involved in stimulus processing, and particularly the PPA) and over-recruitment (i.e., DMN structures, and particularly the Prc), respectively.

In the present study, we used DCM on fMRI data acquired during a visual memory encoding paradigm, in which novel photographs of scenes were presented intermixed with familiar scenes and encoded incidentally via an indoor/outdoor decision task. Memory was tested 70 min later via an old/new recognition memory task with a five-step confidence rating ^8,23,24^ (Figure 1). Successful memory formation was associated with activation of the HC and the PPA as well as deactivation of the Prc in three independent cohorts (cohort 1: 117 young, cohort 2: 58 young, 83 older; cohort 3: 64 young, 84 older; see Table 1). In two cohorts that included both young and older adults, we could further replicate the previously reported age differences in memory encoding with older adults showing reduced activation of the PPA, but relatively increased activity in the Prc (Figure 2), while young and older adults showed comparable involvement of the anterior HC in successful encoding ^5^.

**Figure 1.**
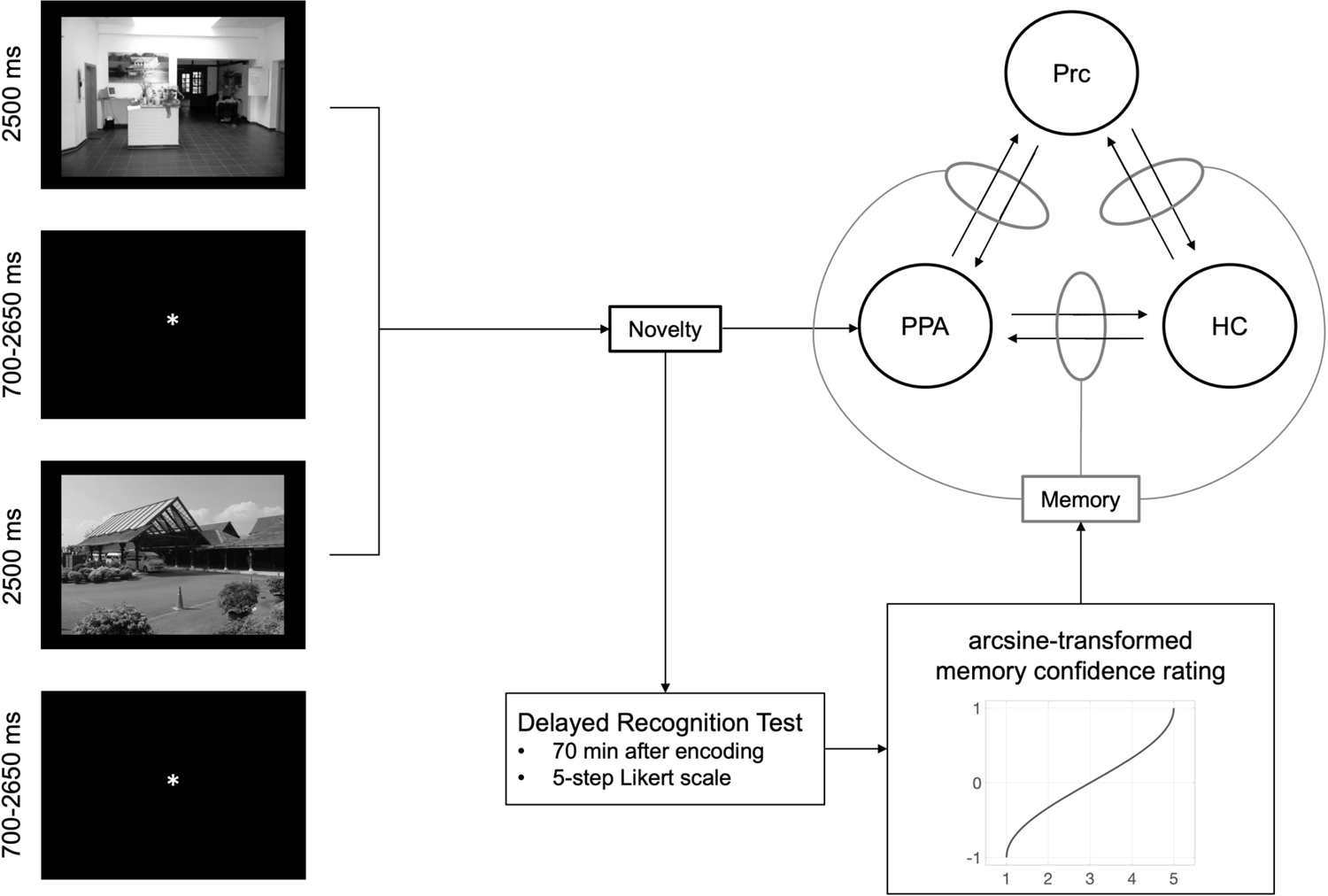
Experimental paradigm and design of the DCM study. During fMRI scanning, participants underwent an incidental visual memory paradigm. We used the novelty response to all novel pictures as driving input to the PPA and generated a parametric memory regressor by parametrically modulating the novelty response with the arcsine-transformed response in the delayed recognition task. The thus obtained memory regressor served as potential contextual modulator of the connections between the PPA, the hippocampus (HC) and the precuneus (Prc).

**Figure 2.**
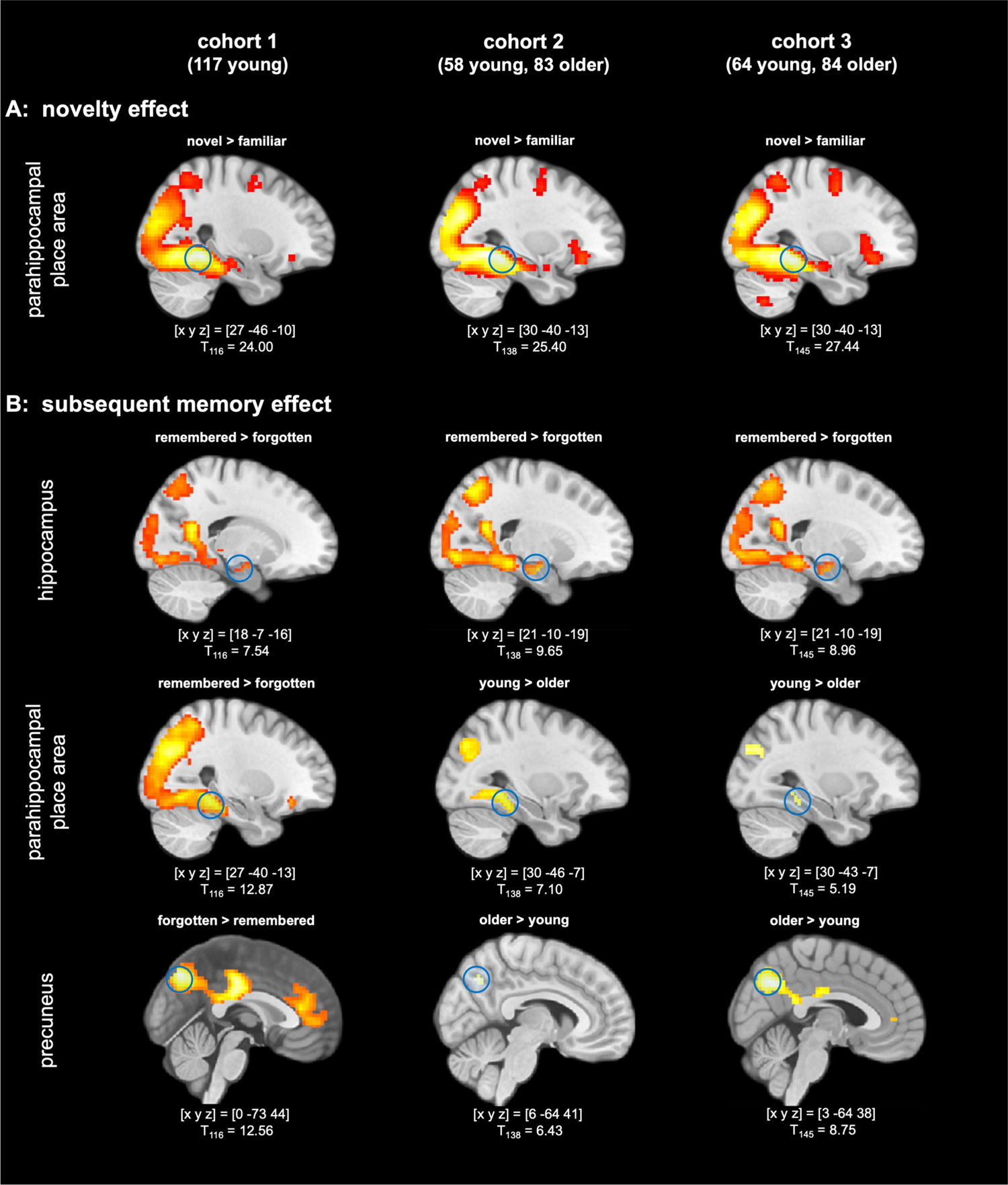
Neural correlates of novelty processing and successful encoding and their age-related differences. **(A)** Novelty effect. Across all cohorts and age groups., novelty processing (i.e., novel > familiar master images) was associated with increased activation of inferior temporo-parietal cortices, particularly the PPA, extending into the hippocampus. **(B)** Subsequent memory effect. Top row: In all three cohorts, successful episodic memory formation elicited activation of the bilateral hippocampus (right hippocampus shown here). Middle row: Encoding-related activation of the PPA (left column) was reduced in older relative to young adults (middle and right column). Bottom row: Encoding-related deactivations in the precuneus (left column) were reduced in older adults (middle and right column). All contrasts are displayed at p < .05, FWE-corrected at voxel-level, minimum cluster size k = 10 adjacent voxels. Coordinates denote local maxima in the respective contrasts.

**Table 1.** Demographic and behavioral data, MRI acquisition parameters.

|  | Cohort 1 | Cohort 2 | Cohort 3 |
| --- | --- | --- | --- |
| **Demographics** |  |  |  |
| *young* |  |  |  |
| N | 117 | 58 | 64 |
| age | **24.37** ($\pm 2.60$) | **23.62** ($\pm 3.45$) | **24.45** ($\pm 4.46$) |
| gender (f/m) | 57 / 60 | 28 / 30 | 37 / 27 |
| *older* |  |  |  |
| N | — | 83 | 84 |
| age | — | **65.20** ($\pm 6.71$) | **62.96** ($\pm 6.06$) |
| gender (f/m) | — | 48 / 35 | 52 / 32 |
| **MRI parameters** |  |  |  |
| MRI scanner | Siemens Verio | Siemens Verio | Siemens Skyra |
| number of slices | 40 | 47 | 47 |
| TR / TE / flip-α | 2.40 s / 30 ms / 80° | 2.58 s / 30 ms / 80° | 2.58 s / 30 ms / 80° |
| voxel size | 2 x 2 x 3 mm | 3.5 x 3.5 x 3.5 mm | 3.5 x 3.5 x 3.5 mm |
| scanning time | 494.4 s | 531.5 s | 531.5 s |
| unwarping | no | yes | yes |
| **Memory performance** |  |  |  |
| *young* A' | **.81** ± .075 | **.83** ± .073 | **.82** ± .068 |
| *older* A' | n/a | **.78** ± .072 | **.77** ± .077 |
| *young vs. older* | n/a | $t_{121.8} = 4.45$;<br>BF<sub>10</sub> = 1417.21 | $t_{142.5} = 3.99$;<br>BF<sub>10</sub> = 203.62 |

Data are shown separately for each cohort. Abbreviations: N = sample size, f = female, m = male, TR = repetition time, TE = echo time, flip-α = flip angle; A' = area under the curve in a ROC analysis of the recognition memory test, reflecting behavioral accuracy (hits vs. false alarms).

Based on these findings, we analyzed patterns of effective connectivity between the hippocampus and temporo-parietal memory network nodes, using the parametric empirical Bayes framework ^25,26^. We constructed a DCM model that included the HC, the PPA and the Prc as regions of interest (ROIs), exposure to novel stimuli as driving input to the PPA, and successful memory formation as a potential parametric modulator at each connection between the three ROIs (Figure 1). The PPA is thought to process and analyze the novel scene, binding its elements and incorporating the current context, which is then bound and encoded together by the HC ^27^. Therefore, we first hypothesized that successful encoding would be associated with an up-regulation of information flow from the PPA to the HC. The Prc constitutes a core structure of a parietal memory network (PMN) proposed by Gilmore, Nelson and McDermott^28^. The PMN has been associated with more pronounced responses to familiar compared to novel stimuli and with negative subsequent memory effects (or “subsequent forgetting effects”) in young adults ^29^, which are attenuated or even inverted in older adults ^5^. Based on those observations, we further hypothesized that successful memory formation would be associated with a down-regulation of information flow (i.e., more pronounced inhibitory effective connectivity) to the Prc from either the PPA or the HC. With respect to age effects, we further hypothesized that at least one of the aforementioned mechanisms – i.e., enhanced information flow to the hippocampus and inhibitory effective connectivity to the Prc - would be attenuated in older adults, which in turn would be associated with poorer memory performance within the group of older adults. To this end, we performed correlational analyses between connection strength and memory performance in the fMRI experiment (area under the curve, A’) ^30^ and in additional, independent tests of explicit memory.

## 2. Results

In the three independent cohorts, a characteristic pattern of novelty-related effective connectivity and encoding-related modulation could be identified. In cohorts 2 and 3, we could further observe age-related reductions in PPA-Prc effective connectivity that correlated with memory performance in the older participants.

### 2.1. Age differences in memory performance

Memory performance was defined as A’, that is, the area under the curve plotting hits (i.e., correctly recognized previously seen images) against false alarms (i.e., new images incorrectly classified as previously seen), taking into account recognition confidence (see methods section for details). Table 1 (bottom) displays A’ separated by cohorts and age groups. In cohorts 2 and 3, we found strong evidence for lower memory performance in older compared to young adults (cohort 2: t_121.8_ = 4.45; BF_10_ = 1417.21; cohort 3: t_142.5_ = 3.99; BF_10_ = 203.62). Gender did not have a robust effect on memory performance in any cohort or age group (all BF_10_ < 0.91), with nominally opposing effects among the older adults of cohorts 2 and 3 (Supplementary Table S1).

### 2.2. Activations and deactivations during novelty processing and memory encoding in young and older adults

Figure 2 displays representative activations and deactivations during novelty processing (novel vs. familiar “master” images) and successful memory encoding (parametric memory regressor). An overview of novelty-related and memory-related fMRI activation maxima, separated by cohorts and age groups, is provided in Supplementary Tables S2-S7.

Novel compared to scene stimuli were associated with a robust fMRI response in the PPA (Figure 2, top row), replicating earlier findings with the same stimulus material ^31^. While PPA novelty responses were stronger in young compared to older adults in cohort 2, this finding could not be replicated at p < .05, FWE-corrected in cohort 3 (Supplementary Tables S3, S4). Regarding the subsequent memory effect, we could replicate previously reported encoding-related activations and deactivations ^4^ in all the cohorts (Supplementary Tables S5-S7). In cohort 1, successful episodic memory encoding was associated with increased activation of the HC (and an extensive temporal and inferior parietal network, including a pronounced local maximum in the PPA (Figure 2, left column). In cohorts 2 and 3, we additionally tested the age-related activation differences during successful encoding (Figure 2, middle and right column; Supplementary Tables S6, S7). Replicating previous studies ^5^, we found older participants to exhibit lower activations in inferior and medial temporal structures, including the PPA, but relatively preserved encoding-related activation of the HC. Furthermore, in line with earlier studies, older adults exhibited reduced deactivations in midline structures of the DMN during successful encoding, with the maximum between-group difference in the right Prc in both cohorts 2 and 3 (Supplementary Tables S6, S7). When explicitly testing for effects of gender, no activation differences survived our statistical threshold of p < .05, FWE-corrected with an extent threshold of 10 voxels in any of the three cohorts.^a^

### 2.3. Locations of regions of interest

Regions of interest (ROIs) of the PPA, the HC, and the Prc were selected based on both structural and functional anatomical constraints (for similar approaches, see ^32,33^). The mean coordinate of the PPA ROI was located near [26-47.5-11], the HC ROI was centered approximately at [26-13.5-17.5], and the mean of the PPA ROI was around [7.3-62.8 38.6] (for an illustration, see Supplementary Figure S1). There was little variability within each cohort (all SD < 3 mm) and high agreement across all cohorts and age groups (Supplementary Table S8).

### 2.4. Temporo-parietal effective connectivity and its modulation during memory encoding

The DCM parameters from the participants of each cohort are displayed in Figure 3, separated by age group for cohorts 2 and 3. Mean DCM parameters are shown in Table 2, separately for each cohort and age group. In every single participant, the driving input (i.e., the PPA response to novel stimuli) was positive (Figure 3, right panel), confirming the plausibility of choosing the PPA as input region.

**Figure 3.**
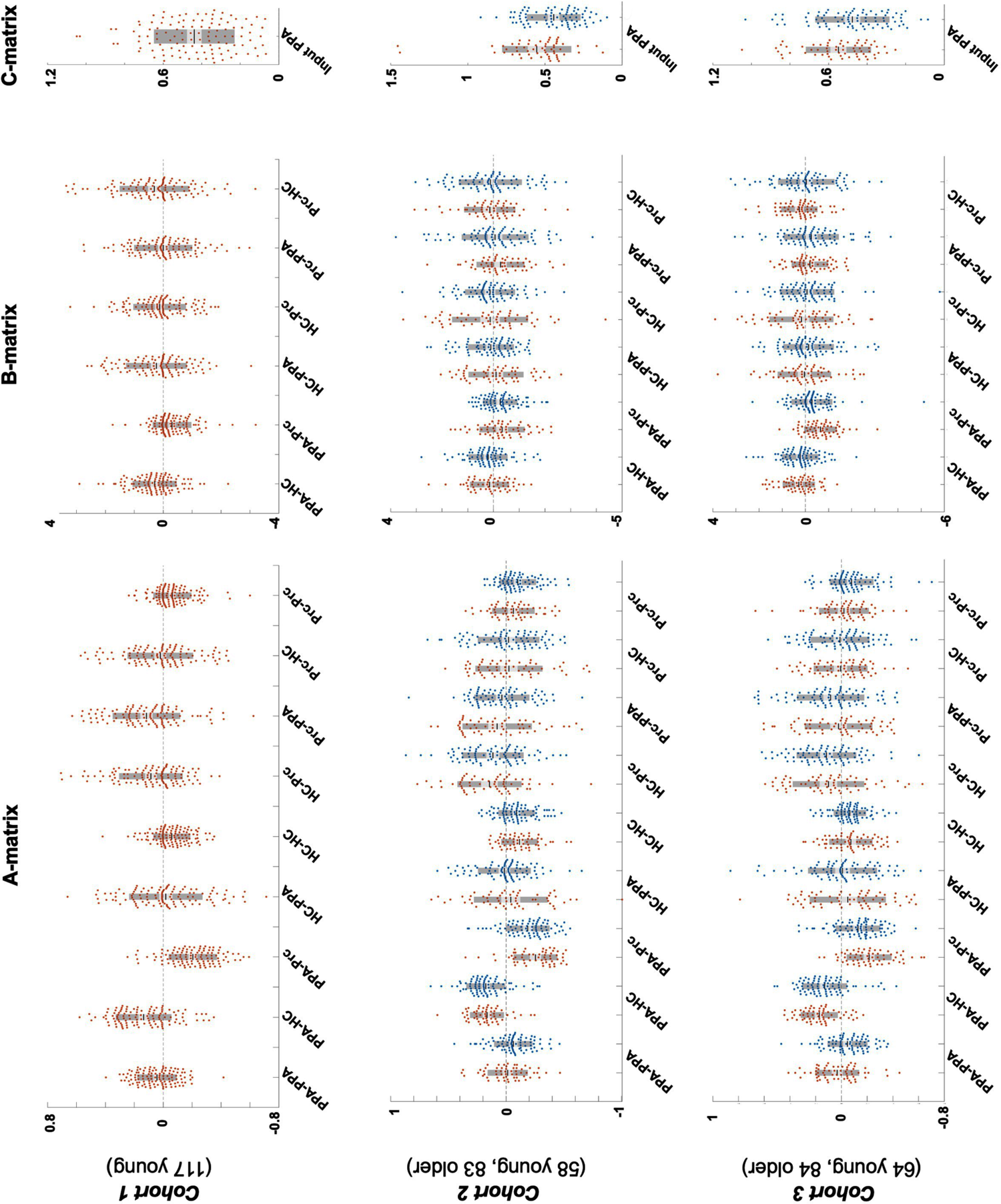
Distribution of DCM parameters across cohorts and single participants. A-matrix: intrinsic connections; B-matrix: contextual modulation by encoding success; C-matrix: driving input by novelty. Plots display means (solid black lines) + / − standard errors (light grey) and standard deviations (dark grey), along with all single data points (red: young adults, blue: older adults).

**Table 2.**
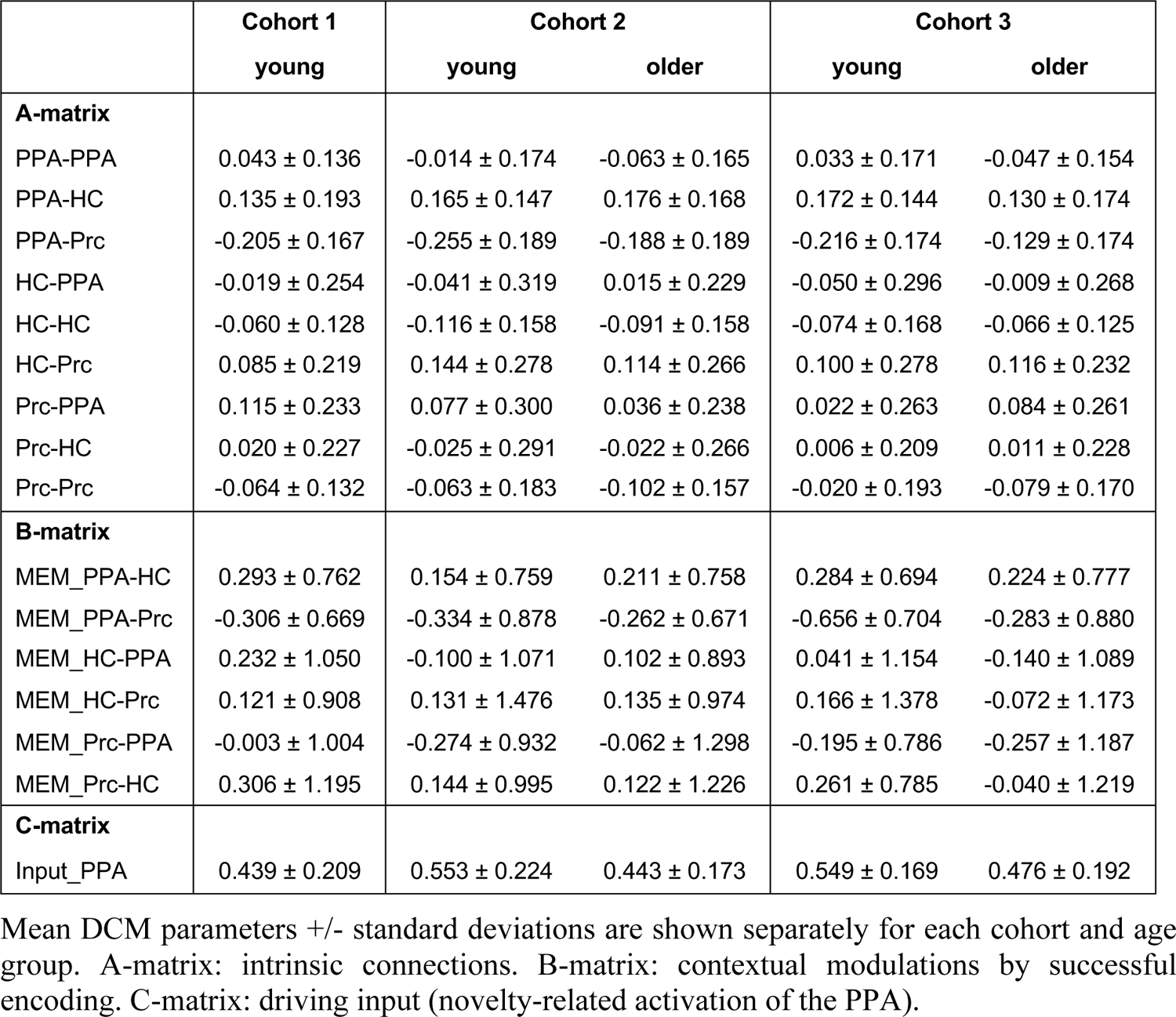
DCM parameters.

Figure 4A displays the patterns of intrinsic connectivity of the temporo-parietal network (A-matrix) during novelty processing investigated in our DCM analyses, separated by main effects (top row) and age differences (bottom row). Connectivity changes related to successful memory formation (B-matrix) are displayed in Figure 4B, following the same layout as Figure 4A. Posterior connection strengths are displayed in Supplementary Figure S2, highlighting connections that could be replicated across all three cohorts (or across cohorts 2 and 3 in case of age group effects).

**Figure 4.**
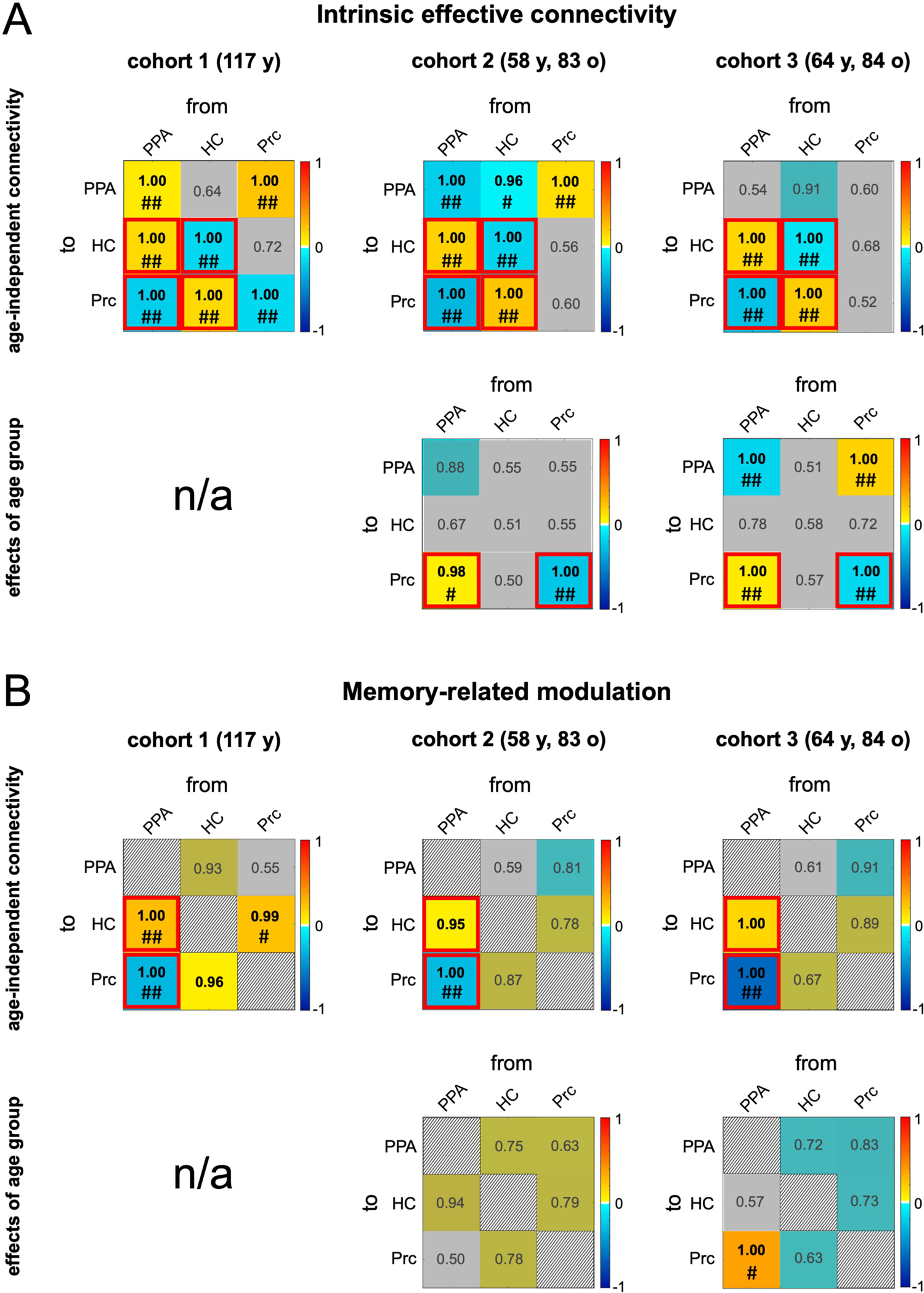
Connectivity matrices of effective connectivity strengths and their modulation by age group. **(A)** Intrinsic connections and **(B)** contextual modulations. Color denotes the parameter size, reflecting positive (“excitatory“) vs. negative (“inhibitory” connectivity for main effects and higher vs. lower values in older adults for age effects. The posterior probability (PP) of a parameter being > 0 in absolute value are shown; connections with PP < .95 are gray-shaded. Hashtags denote free energy-based evidence; ^#^ free energy-based PP > .75 (“positive evidence”); ^##^ free energy-based PP > .95 (“strong evidence”).

#### Intrinsic connectivity between regions

Across all cohorts, there were pronounced excitatory connections from the PPA to the HC and from the HC to the Prc (Figure 3, left panel, Figure 4A, top row). A pronounced inhibitory connection was observed from the PPA to the Prc. All of these connections reached a posterior probability (PP) of 1.00 in all three cohorts (see also Supplementary Figure S2, left panels) and exceeded a PP > .95 also when applying a free energy-based threshold. In cohorts 1 and 2, a positive connection was evident from the Prc to the PPA. In cohort 3, this connection reached a PP of > .95 in older adults only.

#### Intrinsic inhibitory self-connectivity

In the DCM framework, self-connectivity is *a priori* expected to be inhibitory, with higher negative values reflecting stronger (self-)inhibition ^34^. Self-connectivity is reflected in the diagonals of the matrices in Figure 4A (top row). Across all three cohorts, we observed robust negative self-connectivity of the HC (Figure 4A, top row; Supplementary Figure S2, left panels), most likely reflecting reduced auto-inhibition (relative to the implicit baseline) during processing of the novel stimuli, which served as driving input. No replicable pattern emerged with respect to self-connectivity of the PPA or the Prc.

#### Memory-related connectivity changes (contextual modulations)

Successful encoding, as captured by the parametrically modulated memory regressor was associated with increased effective connectivity from the PPA to the HC and more pronounced negative effective connectivity from the PPA to the Prc (Figure 3, middle column, Figure 4B, top row). These connectivity changes could be replicated in all three cohorts at a marginal PP > .95, and the negative PPA-Prc connection also reached a PP > .95 in all three cohorts when applying a free energy-based threshold (Supplementary Figure S2, left panels, red bars). Additionally, in cohort 1, we observed increased bidirectional encoding-related effective connectivity between the HC and the Prc, but, in cohorts 2 and 3, these parameters did not exceed the *a priori* defined threshold of PP > .95.

### 2.5. Age-related difference in memory network effective connectivity

As there were no older adults in cohort 1, all age differences reported here are based on cohorts 2 and 3. In both cohorts, age group was associated with more positive (i.e., reduced inhibitory) effective connectivity from the PPA to the Prc (Figure 4A/B, bottom rows; Supplementary Figure S2, right). This reduced inhibitory temporo-parietal effective connectivity was found in the A-matrix (intrinsic connection) in both cohorts, and, additionally, in cohort 3, it was also found in the B-matrix (memory-related contextual modulations). Furthermore, even though there was no clear age-independent pattern (see 3.2) of Prc self-connectvity across the three cohorts, inhibitory Prc self-connectivity was nevertheless reduced in older adults compared to young adults in both cohorts 2 and 3.

### 2.6. Association of effective connectivity with memory performance in older adults

As described above, the inhibitory connection from the PPA to the Prc was attenuated in older compared to young adults (Figure 4A, bottom row; Supplementary Figure S2, right panels). We therefore focused on this connection when testing for a relationship between DCM parameters and memory performance in older adults. Outlier-robust Shepherd’s *Pi* correlations ^35^, adapted for Bayesian statistics ^36^, were computed between the PPA-Prc connection strength and A’ as measure of memory performance (see Methods section for details). In both cohorts (2 and 3), the parameter representing PPA-Prc connectivity was negatively correlated with A’ (cohort 2: ΙΙ = −0.368, BF_10_ = 18.21; cohort 3: ΙΙ = −0.297, BF_10_ = 2.63; Figure 5A), such that stronger PPA-Prc inhibitory connectivity related to better memory performance.

**Figure 5.**
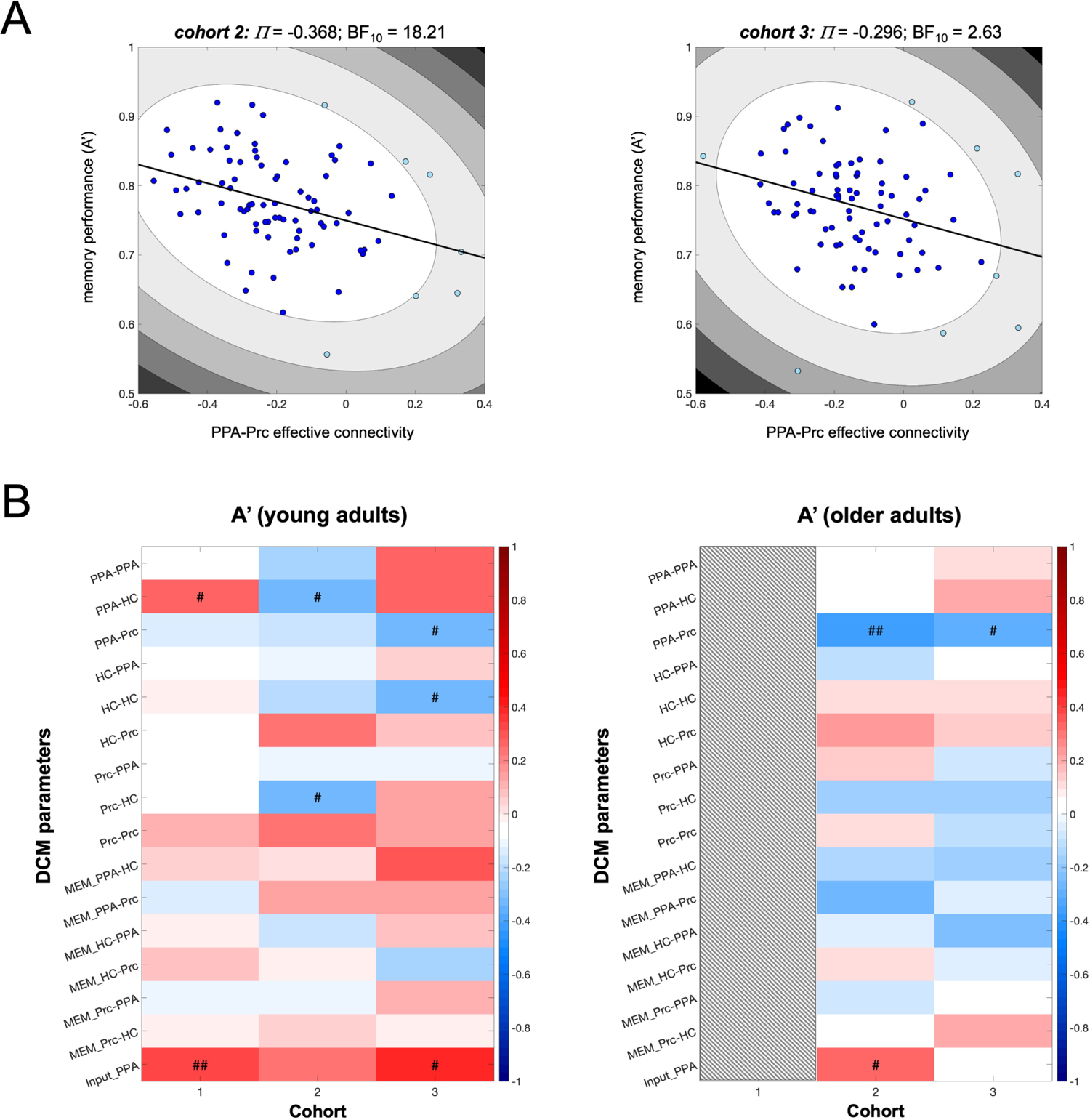
Correlations between DCM parameters and memory performance. **(A)** Plots depict outlier-robust Bayesian Shepherd’s Pi correlations between the inhibitory PPA-Prc connection in older adults in cohorts 2 and 3. **(B)** Results from pair-wise Bayesian Shepherd’s *Pi* correlations of the individual participants’ DCM parameters with memory performance (A’) in young (left) and older (right) participants, separated by cohort. Positive and negative correlations are shown in red and blue, respectively. ^#^ BF_10_ > 2.00; ^##^ BF_10_ > 10.00.

We then computed exploratory Bayesian Shepherd’s *Pi* correlations for A’ and all DCM parameters in young and older adults (Figure 5B). In older adults of both cohorts, we found replicable evidence only for the correlation with the PPA-Prc connection. In young adults, evidence in favor of a correlation between the PPA-Prc connection and A’ was found in cohort 3 (ΙΙ = −0.317, BF_10_ = 2.01), but not in cohorts 1 and 2 (all BF_10_ < 0.23). On the other hand, the driving input (C-matrix, i.e., the PPA novelty response) was correlated positively with A’ in young adults in cohorts 1 and 3, but not in older adults. Calculating cumulative Bayes factors as the Bayes factors for the averaged correlation coefficients (weighted for effective sample sizes)^b^, confirmed the observation of correlations between the driving input and A’ in young adults and between the PPA-Prc connection and A’ in older adults (all cumulative BF_10_ > 400.0), The cumulative evidence for a relationship between the PPA-Prc connection in young adults, on the other hand, was limited (BF_10_ = 3.96; Supplementary Tables S9, S10).

Finally, we tested for a potential relationship between DCM parameters and measures of memory in independent – verbal – memory tests, the Verbal Learning and Memory Test (VLMT) ^37^ and the Logical Memory subscale of the Wechsler Memory Scale (WMS) ^38^. No correlations of DCM parameters with those measures of memory performance were directly replicable across cohorts for either age group. In the cumulative statistics there was evidence for a relationship between VLMT and WMS performance (learning and delayed recall) and the driving input in older, but not in young adults, but the evidence was substantially lower (15.0 < BF_10_ < 19.8) Supplementary Figure S3, Supplementary Tables S9, S10).

## 3. Discussion

In the present study, we have employed DCM to elucidate the putative temporo-parietal information flow underlying fMRI activations and deactivations during successful memory formation and their characteristic age-related differences. We found that successful memory encoding was associated with increased PPA-hippocampal effective connectivity and increased inhibitory effective connectivity from the PPA to the Prc in both young and older adults. In older adults, the inhibitory connection from PPA to Prc was, however, reduced, and less negative inhibitory – or even positive – PPA-Prc connectivity was associated with poorer memory performance.

### 3.1. Cortical-hippocampal interactions during novelty processing and memory encoding

In three independent cohorts, a characteristic pattern of intrinsic effective connections emerged, which were in part further modulated as a function of encoding success (for an overview, see Figure 6). Most prominently, the following connections were observed:

1. *The PPA exerts novelty-related excitatory input on the HC, which is up-regulated during successful encoding*. Previous studies have demonstrated that both the HC and the PPA exhibit robust responses to novel scenes ^39,40^ as well as subsequent memory effects ^30,41^. The direction of information flow from the PPA to the HC observed in the present study suggests that HC-dependent encoding of the stimuli occurs *after* higher-level perceptual analysis of the scene stimuli in the PPA. This interpretation is supported by previous neuroimaging studies ^27,42^ as well as by electrophysiological investigations and immediate-early gene imaging studies in rodents, which, despite the lack of a clearly defined homologue of the PPA in rodents, support the notion that scene processing is mediated by the parahippocampal cortex ^43^ and that hippocampal encoding of episodes is preceded by stimulus processing in inferior-medial temporal cortices ^44,45^.
2. *The HC shows negative self-connectivity during novelty processing.* Self-connectivity is *a priori* defined as negative within the DCM framework ^34^, with negative self-connectivity reflecting the release of auto-inhibition. Negative self-connectivity of the HC upon driving input (here: activation of the PPA to novel scene stimuli) ^46^, in our view, thus most likely reflects hippocampal novelty responses to the scene stimuli. The HC exhibits strong responses to novel stimuli ^47,48^, a finding also replicated in another cohort of older adults using the same paradigm as in the present study ^49,50^. Hippocampal self-connectivity was not influenced by age group in our study. Age-related changes in intra-hippocampal connectivity have been reported using different methodologies, although results are not entirely conclusive. While a DCM study on navigational learning found reduced hippocampal auto-inhibition in older adults ^22^, results from longitudinal resting-state fMRI investigations suggest an age-dependent decrease of voxel-wise functional connectivity in the anterior HC, but a connectivity increase in the posterior HC ^51^. A limitation of our present study is that we did not allow for a modulation of self-connections by successful encoding in our model, as the self-connections of the PPA and Prc did not replicate consistently across cohorts (Figure 4). We can therefore not exclude a potential age-dependent modulation of hippocampal self-connectivity during successful encoding.
3. *The PPA exerts inhibitory input to the Prc during novelty processing and successful encoding.* Deactivations of the Prc are a common finding in studies employing the subsequent memory effect ^4,29^, and a straightforward explanation for this is that the Prc as a DMN structure deactivates during most tasks that require externally directed attention ^52^, including novelty processing ^50^ and successful encoding ^5,24^. During encoding of novel stimuli, it may be of particular importance to suppress processing related to retrieval and familiarity, which engages the Prc ^4,28,53^. While the function of the Prc in explicit memory processes likely goes beyond supporting retrieval (see 4.2), it is notable from the present results that suppression of Prc activity during novel stimulus processing and successful encoding was not found to be mediated by the HC, but rather directly by the PPA, bypassing the HC.
4. *The HC exerts excitatory input to the Prc during novelty processing.* Unlike the PPA, the HC exerted positive input to the Prc during processing of novel stimuli, although there was only very limited evidence for a further modulation by encoding success (Figure 3, 4, Supplementary Figure S2). One explanation for this may be that the hippocampal novelty signal is relayed to the Prc, allowing for novelty-related deactivation ^28^ (Section 4.2).
5. *The Prc exhibits positive effective connectivity to the PPA during novelty processing.* In cohorts 1 and 2 and in the older participants from cohort 3, we further found an excitatory connection from the Prc to the PPA. The Prc is strongly interconnected with the parahippocampal cortex ^54^, of which the PPA is a subregion. Projections from the Prc to the parahippocampal cortex are thought to be of a feed-backward type. One possible function of this connection could be that the Prc provides information based on existing knowledge that can be used to integrate elements of a novel scene into existing memory representations (for a detailed discussion, see 4.2).

**Figure 6.**
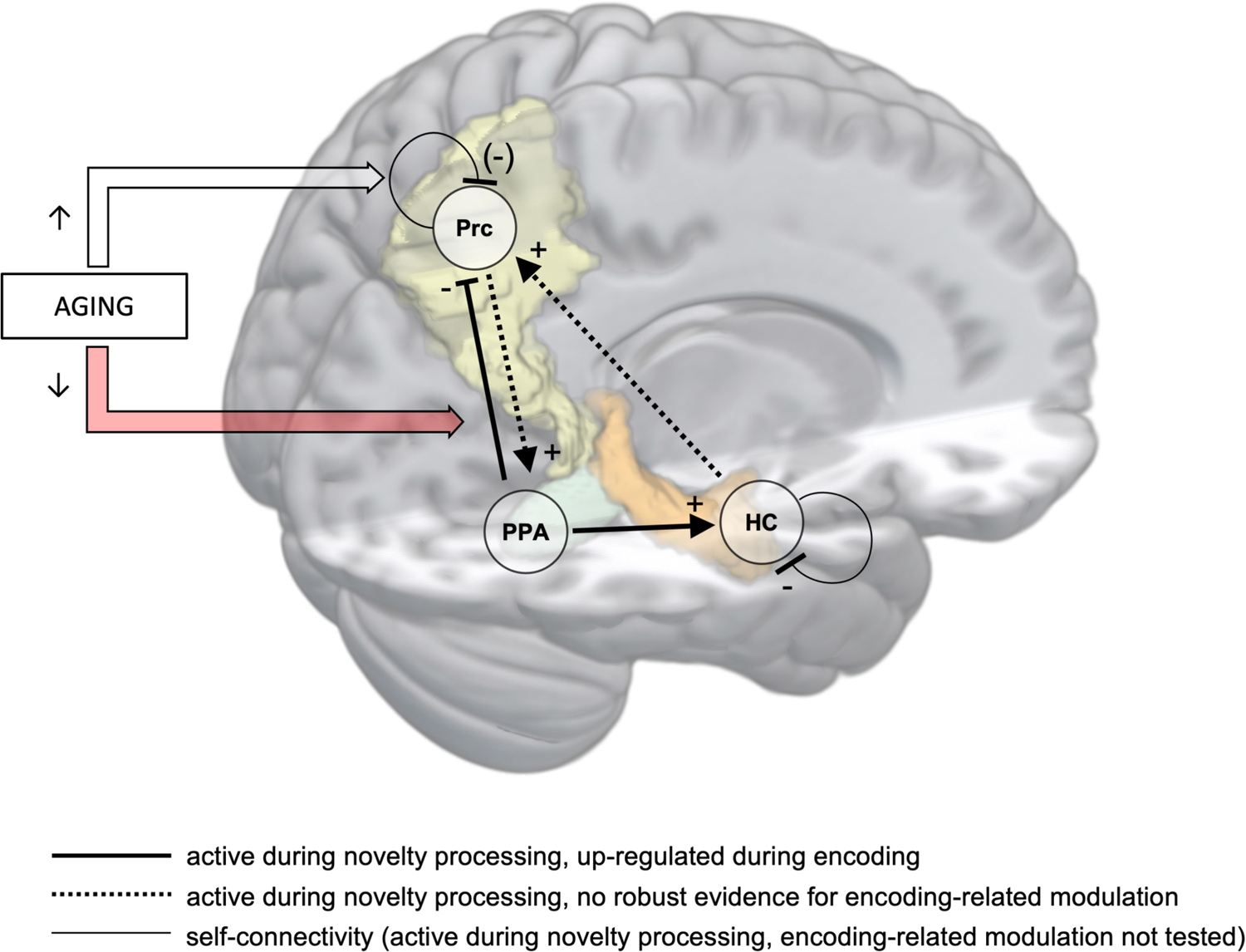
Temporo-parietal effective connectivity during novelty processing and successful memory encoding. Successful encoding was associated with increased activation of the hippocampus by the PPA and stronger suppression of the precuneus by the PPA. In older adults, the inhibitory connection from PPA to precuneus was attenuated and related to worse memory performance.

### 3.2. Reduced memory-related deactivations of the precuneus in older adults

The DMN, and particularly the Prc typically shows activations during episodic retrieval, but deactivations during encoding of novel stimuli ^4^ (for a potentially differential role of the Prc during encoding of repeatedly presented information, see Supplementary Discussion). It thus seems plausible to assume that, during processing of novel stimuli, suppressing Prc activity may help to reduce interference with familiar information ^28^. The results of our DCM analysis point to the presence of a direct inhibition of the Prc by the PPA during processing of novel stimuli, which is further enhanced during successful encoding and attenuated in older compared to young adults (Figure 6). We therefore tentatively suggest that the commonly observed attenuated encoding-related DMN deactivation in older adults – and especially in those with poor memory performance – ^5,24^ may result from a reduced ability to suppress ongoing DMN activity during novelty processing and successful encoding.

This interpretation may be somewhat at odds with the commonly proposed notion that increased activation of parietal – and also prefrontal – neocortical structures may reflect a compensatory mechanism for diminished MTL function in old age ^5,55^. One possible interpretation for this apparent discrepancy is that compensatory activity might be particularly relevant in older adults with pre-clinical or subclinical memory impairment. While preserved patterns of memory-related brain activity (i.e., patterns with high similarity to those of young adults) have been associated with better memory performance in older adults ^8,30,56,57^, those with poorer memory performance can show not only deactivations, but even atypical activations of DMN structures during successful versus unsuccessful encoding ^5^, possibly reflecting an increased reliance on self-referential or prior knowledge-dependent information during encoding ^58–60^. In line with this interpretation, increased activity of midline brain structures during novelty processing and successful memory encoding has been reported in individuals at risk for Alzheimer’s disease (AD), that is, otherwise healthy older adults with subjective cognitive decline ^50^ or cognitively unimpaired older individuals with increased beta-amyloid deposition as assessed with PET ^61^. Notably, this DMN hyperactivation is no longer observed in patients with early-stage AD ^50^, suggesting that, whether compensatory or not, it is nevertheless vulnerable to more pronounced disruption of brain function. Whether DMN hyperactivation reflects compensatory mechanisms at a pre-clinical stage that break down during disease progression or rather a more general impairment of inhibitory processes that precedes a broader decline in neural function should be further elucidated in future studies.

### 3.3. The relationship between inhibitory connectivity and memory performance

In our present study, only the inhibitory connection from the PPA to the Prc was associated with memory performance in older adults, whereas evidence for an association of that connection with memory performance in young adults was limited (Figure 5, Supplementary Tables S9, S10). This suggests that neocortical inhibitory mechanisms, in addition to altered hippocampal connectivity ^21^ contribute to age-related reduced efficiency of memory networks. This does not rule out that connectivity of the hippocampus with other (e.g., prefrontal) brain regions may also be associated with memory performance. Nyberg and colleagues, for example showed that higher connectivity of the ventrolateral PFC and the hippocampus connectivity during encoding was associated with higher dropout rate from a longitudinal study, as a proxy for worse cognitive and clinical outcome ^21^. Furthermore, a study on hippocampal-posterior neocortical effective connectivity during spatial navigation suggested that reduced hippocampal auto-inhibition was associated with lower learning performance in older adults ^22^. Despite the heterogeneity of the models and ROIs across those studies and our present study, we suggest that our findings nevertheless converge with the aforementioned results in highlighting, more generally, the role of inhibitory connectivity within the extended memory networks in successful aging of human memory systems.

It should further be noted that age-related activation increases in brain structures not typically involved in the task at hand have also been observed in cognitive domains other than explicit memory ^62^, and even in the motor system ^63^. As, in the latter study, ipsilateral motor system activity did not predict better performance (e.g., reaction times), the authors suggested that their results speak against a compensatory role of age-related task-related hyperactivations. Together with our present results, we suggest that loss of inhibitory activity might constitute a cross-domain mechanism of brain aging, and future research should further explore its possible relationship with subclinical pathology.

### 3.4. The multifaceted role of the precuneus in human long-term memory

The engagement of DMN structures during memory *retrieval* processes is a well-replicated finding ^52,53^, and the Prc, in particular, is well-known for its role in episodic retrieval ^4^, despite an ongoing debate with respect to a preferential role in recollection versus familiarity ^28,64^. Irrespective of the type of retrieval process involved, it seems plausible to assume that, during processing of novel stimuli, suppressing Prc activity may help to reduce interference with familiar information. In line with this interpretation, Gilmore and colleagues proposed a parietal memory network that includes the Prc, which deactivates in response to novel information and shows increased responses as a function of stimulus familiarity ^28^. During successful encoding, higher activity in ventral parietal structures has been associated with subsequent forgetting rather than remembering ^4,28,29^. On the other hand, there has also been evidence for higher activity in dorsal parietal structures associated with positive subsequent memory effects ^29^. While the meta-analysis by Uncapher and Wagner was focused on lateral parietal structures, a similar pattern of functional heterogeneity has also been proposed for midline parietal subregions of the DMN ^65^. Particularly the dorsal Prc is involved in switching between DMN activity and activity of task-positive networks ^66^ and has been shown to be actively engaged in some attention-demanding tasks, possibly mediating the integration of inwardly-directed and externally-directed cognitive processes ^67^. More recently, the precuneus has also been proposed as a key region involved memory acquisition, particularly for visuo-spatial material ^68–70^. However, those studies all employed repeated stimulus exposure, and may therefore reflect a primarily neocortically-mediated form of learning distinct from hippocampus-dependent memory for unique episodes ^71,72^.

One possibility to reconcile those findings with the results of our present study is that the Prc may serve, to some extent, as a gatekeeper during long-term memory formation, enabling the parallel storage of unique episodes in their spatial and temporal context and of elaborate and multiply associated information (i.e., schemas ^71,73^. More broadly, future research should explore the possibility that parietal memory network structures may contribute to the distinction of episodic and semantic memory ^74^.

### 3.5. Associations of the driving input with memory performance

While, in young adults, we found only limited evidence for association between PPA-Prc inhibitory connectivity and memory performance, the combined analyses across cohorts point to very strong evidence for a positive relationship between the driving input to the PPA and memory performance in young adults (A’; see Supplementary Table S10). In older adults, such a relationship was not found for A’, but there was moderate evidence for a positive association between learning and delayed recall performance in the WMS and, to some extent, also in the VLMT (Supplementary Table S9).

With respect to our present results, we tentatively suggest that the most straightforward observation for the correlations observed here is that C-matrix in our DCM design essentially reflects the novelty response in the PPA. Previous analyses of our dataset suggest that a higher similarity of the whole-brain novelty and memory responses with the prototypical responses in a reference sample of healthy young adults is associated with better memory performance in both the experimental task itself and also in the WMS and VLMT ^30,57^. There were a few differences in details between the correlations here and those reported in our previous works (e.g., overall stronger correlations of the memory-based scores with memory performance, particularly in younger adults), and we suggest that they most likely reflect the definition of the underlying fMRI measures (whole-brain single-value scores vs. the peak novelty response in the PPA), which, despite likely showing high covariance, nevertheless constitute distinct measures of brain activity. According to this interpretation, the observed correlations between the driving input and the performance in independent memory tests in older adults may reflect the previously described associations between the overall integrity of fMRI response patterns and cognitive function, particularly memory performance ^56,57^.

### 3.6. Feasibility and limitations of replication in DCM studies

Beyond the specific implications for cognitive neuroscience of the aging memory system, our present study also concerns the critical issue of replication in human fMRI research ^75^. While recent analyses suggest that the replicability of group-level fMRI results is overall encouraging when a study is adequately powered ^76,77^, few studies have so far addressed the issue of replication in DCM, and even less so, in the context of the parametric empirical Bayes (PEB) framework employed here. Earlier DCM study describing replications or addressing the issue of reproducibility have commonly reported results from Bayesian model selection in combination classical statistical tests (e.g., t-tests) on the individual participants’ DCM parameters ^78–80^ or even fully relied on classical statistical testing at group level ^81^. In the present study, we observed highly reproducible patterns of intrinsic connectivity in four parameters of the A-matrix (intrinsic connectivity during novelty processing; see Figure 4A, top row), all of which remained in the model when applying the more stringent free energy-based threshold. On the other hand, self-connections, particularly those of the PPA, showed considerable inconsistencies, even with robust effects in opposite directions across cohorts (Figure 4A). This observation points to a potentially underappreciated difficulty in robustly modeling self-connections, which might also explain the rather low rate of reporting self-connectivity in neuropsychiatric DCM studies (approximately 25%) ^82^. Thus, we did not include a modulation of self-connectivity in our model space, as we found it difficult to generate hypotheses on the modulation of self-connections that showed no robust intrinsic pattern.

In the B-matrix (contextual modulation by successful encoding), the enhanced negativity of the PPA-Prc connection could be replicated at free energy-based PP > .95 (Figure 4B, top row). The increased positivity of the PPA-HC connection was robust at PP > .95 (free energy-based) in cohort 1 (which consisted of young participants only), whereas this connection was reliable only as marginal PP in cohorts 2 and 3 (which included approximately 57% older adults). One reason for this may be that the free energy threshold is based on iterative pruning of parameters from the model based on the posterior covariance among parameters ^26^, This procedure is highly effective for the selection of the most parsimonious models, but, in the context of replication, one should consider the possibility that subtle differences in the strengths of parameters with high covariance across cohorts may result in differential pruning, which might, in extreme cases, result in an apparent replication failure when considering free energy based thresholds only. Between-group differences might also explain the inconclusive findings regarding the bidirectional positive modulation of HC-Prc connectivity observed in cohort 1. Despite showing nominally positive parameters in cohorts 2 and 3, these connections did not pass the *a priori* defined threshold of PP > .95 in these cohorts (Figure 4B). In addition to the inclusion of older participants, the differences in trial timings and scanning parameters (Table 1) might constitute factors that could potentially impair robust replication.

In addition to the DCM parameters themselves, replicability is also important with respect to their correlation with behavioral measures. While we found negative correlations between PPA-Prc connectivity and memory performance in the older adults of both cohorts 2 and 3, the strength of evidence between the cohort was different, despite similar magnitudes of the correlation coefficients (Figure 5A). Here we aimed to implement a meta-analytic approach to obtain cumulative Bayesian inference across cohorts (Nikolakopoulos and Ntzoufras, 2021; see Supplementary Tables S9, S10), but we acknowledge that there is an ongoing methodological debate on cumulative Bayesian inference ^83–85^ and that further validation of the method used here is warranted. This is of particular importance with respect to the correlations with independent measures of memory, for which evidence was found only in the cumulative statistics (Supplementary Table S9). Furthermore, additional replication approaches to assess the robustness of brain-behavior correlations in DCM, such as bootstrapping ^86^ should be considered in future studies.

### 3.7. Interpretation of DCM results at the neuronal level

While the results of our DCM analyses could be robustly replicated in three independent cohorts, it must nevertheless be acknowledged that DCM approach relies on multiple highly complex assumptions regarding functional neuroanatomy (see Section 3.5 in the main manuscript), model space and model selection ^87,88^, as well as the underlying biophysical mechanisms when inferring on a neuronal state from a hemodynamic signal ^88^. In the present study, we interpreted our main finding, such that the PPA exerts inhibitory activity onto the precuneus during novelty processing, which is further increased during successful episodic encoding and attenuated in aging. More generally, we have used DCM to infer on information flow between brain areas in relation to a driving input (novelty) and a behavioral outcome (remembered vs. forgotten) contextually modulating connections between regions. Information flow in this context can be thought of as the first region passing on information to the second region (by causing it to receive different inputs as a function of the contextual modulator). Similarly, a region can be thought of as acting inhibitory or excitatory on a target area, if the strength of its connection to this area is negative or positive, respectively, in line with widely accepted interpretations of DCM ^88^.

Thus, it must be noted that, when referring to inhibitory activity in the interpretation of our present results, we refer to large-scale network dynamics, not small-scale regional pertubations. We acknowledge that great care must be taken when interpreting the results of DCM for fMRI data with respect to the underlying neuronal processes, particularly at the microcircuit level. One promising approach for a future improvement is to employ more elaborate models of the underlying neurovascular coupling and resulting BOLD signal ^89–91^, which can, in principle, be incorporated into DCM analysis ^92^.

### 3.8. Conclusions

Using dynamic causal modeling on fMRI data acquired during a visual memory encoding task, we could demonstrate that successful memory formation is associated with increased information flow from the PPA to the hippocampus and suppression of information flow from the PPA to the precuneus. A preserved inhibitory PPA-precuneus connection was associated with better memory performance in two cohorts of healthy older adults, suggesting that, in addition to age-related changes in hippocampal connectivity, neocortical connectivity changes should also be considered. Our results highlight the importance of inhibitory mechanisms in successful aging of the human memory system. Future studies should investigate the longitudinal trajectories of inhibitory processes in human explicit memory, in order to explore the possibility that impaired inhibition might constitute an early pathophysiological mechanism in the development of age-related cognitive decline.

## 4. Limitations

### 4.1. Regions of interest and the comparability of DCM studies

A limitation of the present study is inherent to the DCM approach, that is, the restriction to the *a priori* selected regions included in the model ^87^. In the present study, we performed a rather rigid ROIs definition, based on anatomical constraints, literature-based assumptions, and fMRI activations, similarly to a previously described approach ^33^ (see Methods, 2.6.1). While this approach yielded highly consistent ROI locations across cohorts and age groups (Supplementary Table S8), the choice of ROIs nevertheless differed from those used in previous DCM studies ^21,22,93^. Most critically, our model did not include prefrontal cortex regions, which are typically involved in successful episodic encoding ^4^. One reason for this was that the focus of our study was on the Prc, which showed the most robust age-related activation difference (Figure 2; Supplementary Tables S6, S7). Furthermore, encoding-related prefrontal activation clusters were distributed across several frontal lobe structures and showed variability across cohorts, such that selection of one given prefrontal ROI might have been to some extent arbitrary. We can therefore not exclude that prefrontal connectivity may additionally contribute to memory performance older adults, as suggested in earlier studies ^21,94^.

Within the hippocampus, we restricted our ROI to the anterior HC. This was motivated by previously reported associations between anterior hippocampal structure and function with age-related memory decline ^21,95^ and on the aim to avoid an overlap with the PPA ROI, but we acknowledge that investigating differential connectivity patterns along the hippocampal axis will likely provide further important information on the aging hippocampus-dependent memory system ^96^.

Another limitation related to our ROI selection concerns the choice of the PPA as input region. While it is plausible to employ either early visual cortex regions ^21,22,93^ or stimulus-responsive brain structures (e.g., the fusiform face area in a face-name encoding task ^21,22,93^), this may limit the generalizability of the resulting models to other cognitive tasks. In the present study, for example the observed association between PPA-Prc connectivity and memory performance did not generalize to independent, verbal, memory tasks (Supplementary Figure S3, Supplementary Table S9), which suggests that the specific association of PPA-Prc effective connectivity with memory performance may be limited to visual or spatial memory.^c^

### 4.2. Further limitations and directions for future research

Beyond ROI selection, it must be noted that the DCM approach relies on complex assumptions with respect to functional neuroanatomy (see above), model space and model selection ^87,88^, but also the underlying biophysical mechanisms when inferring on a neuronal state from a hemodynamic signal ^88^. It is thus likely overstated to assume direct causality from the effective connectivity patterns observed, and we are aware that our model, its reproducibility aside, is most likely incomplete (for a more detailed elaboration, see Supplementary Discussion 1.3).

Another limitation of the present study may result from the sampling of the study cohort ^30^ and the lack of AD-related biomarkers. Despite the considerable interindividual variability in memory performance among the older adults in our cohort (^24^, Figure 2), it must be acknowledged that older participants were most likely healthier than average. This was both due to our participant recruitment strategy, which led to healthy and active older adults to more likely volunteer for participation (see Methods, 2.1), and due to the exclusion of participants with severe comorbidity ^30^. In a large cohort that included older adults with subjective cognitive decline (SCD) and mild cognitive impairment (MCI) as well as manifest AD, novelty-related Prc responses were found to exhibit an inverse U-shaped pattern from healthy older adults over SCD and MCI to manifest AD ^50^. It remains thus to be determined whether the association between temporo-parietal inhibitory connectivity and memory performance also applies to individuals with memory impairment or dementia risk states. Furthermore, future studies should also follow up older adults with memory impairment longitudinally to assess whether progressive memory decline might be associated with decreasing inhibitory connectivity over time.

## 5. Notes

### 5.1. Author contributions

Conceptualization: B.H.S., J.S., G.Z.; Investigation: A.A., A.R.; Methodology: B.H.S., J.S.; Formal analysis: B.H.S., J.S., A.R.; Project administration: B.H.S., A.R.; Software: B.H.S., J.S., H.S.; Supervision: B.H.S.; Visualization: B.H.S.; Writing – original draft: B.H.S.; Writing – editing: B.H.S., J.S., J.M.K., A.M., G.Z., M.S., A.R.; Funding acquisition: B.H.S. All authors approved the final version of the manuscript.

## Acknowledgments

The authors would like to thank Nadia Ay, Hannah Feldhoff, Larissa Fischer, Sophia Jauch, Lea Knopf, Matthias Raschick, Marc Roder, and Annika Schult for help with neuropsychological data collection and analysis and Kerstin Möhring, Katja Neumann, Ilona Wiedenhöft, and Claus Tempelmann for assistance with MRI data acquisition. We further thank Emrah Düzel for helpful discussion during the planning stage and collaboration during data collection.

## 5.2. Funding Statement

This study was supported by the State of Saxony-Anhalt and the European Regional Development Fund (Research Alliance “Autonomy in Old Age” to B.H.S.) and by the Deutsche Forschungsgemeinschaft (DFG CRC 1436, TP A05 to B.H.S., TP B01 to M.S., TP C01 to G.Z., and TP Z03 to A.M.; DFG RI 2964-1 to A.R.). The funding agencies had no role in the design or analysis of the study.

## 5.3. Inclusion and Diversity

We support inclusive, diverse, and equitable conduct of research.

## 5.4. Data Availability Statement

Matlab scripts used for data analysis are available via GitHub (https://github.com/bhschott/DCM_MemoryEncoding). Group GLM fMRI results (novelty and memory contrasts, separated by cohort and age group) and group DCM files (GCM*.mat) are available via OSF (https://osf.io/ktqy6/). Due to restrictions imposed by the local ethics committee and EU data protection regulations, sharing of the single subjects’ (f)MRI data sets underlying this study in a public repository is not possible. Access to de-identified raw data will be provided by the authors upon reasonable request. Please contact the corresponding author (B.H.S.) for data requests.

## 8. Methods

### Key resources table

| REAGENT or RESOURCE | SOURCE | IDENTIFIER |
| --- | --- | --- |
| Deposited data |  |  |
| Collated single subjects' DCMs and templates (GCM_*.mat) | Björn H. Schott via OSF | https://osf.io/ktqy6/ |
| Group level fMRI contrasts | NeuroVault | https://osf.io/ktqy6/ |
| ROI masks (PPA probabilistic ROI and anatomical masks for all ROIs) | Zweynert et al., 2011 / Björn H. Schott via OSF | https://osf.io/ktqy6/ |
| Single subjects' raw fMRI data (Autonomy in Old Age) | Leibniz Institute for Neurobiology (LIN) | n/a |
| Software and algorithms |  |  |
| Matlab R2018b | MathWorks, Natick, MA | https://de.mathworks.com/products/matlab.html |
| Statistical Parametric Mapping (SPM12, v7771), including DCM 12.5 | Wellcome Trust Centre for Neuroimaging | https://www.fil.ion.ucl.ac.uk/spm/ |
| Parametric first-level fMRI data analysis described by Soch et al., 2021<sup>23</sup> | Joram Soch via GitHub | https://github.com/JoramSoch/FADE_BMS |
| Matlab scripts for ROI selection, DCM batch analysis, and brain-behavior correlations | Björn H. Schott via GitHub | https://github.com/bhschott/DCM_MemoryEncoding |
| WFU Pickatlas | Wake Forest University | https://www.nitrc.org/projects/wfu_pickatlas/ |
| Shepherd's *Pi* Correlation | Sam Schwarzkopf | https://www.frontiersin.org/articles/10.3389/fnhum.2012.00200/full |
| Matlab Bayes Factor Toolbox | Bart Krekelberg via GitHub | https://github.com/klabhub/bayesFactor |
| UnivarScatter plot function | Manuel Lera Ramirez via GitHub | https://github.com/manulera/UnivarScatter |

### Resource availability

#### Lead Contact

Further information and requests for resources should be directed to the lead contact, PD Dr. Dr. Björn H. Schott.

#### Materials availability

This study did not generate plasmids, mouse lines, or unique reagents.

#### Data and code availability

*Code:* The code generated in the course of data analysis is available from the lead contact’s GitHub site (https://github.com/bhschott/DCM_MemoryEncoding).

*Data:* DCM results reported in the present study and the corresponding group-level fMRI results (novelty and subsequent memory contrasts) are available from OSF (https://osf.io/ktqy6/). The single subjects’ raw MRI data reported in this study cannot be deposited in a public repository due to restrictions imposed by the local ethics committee and EU data protection regulations. To request access, contact please contact the lead contact.

Any additional information required to reanalyze the data reported in this paper is available from the lead contact upon request.

### Experimental model and study participant details

#### Participants

In the present study, we investigated three cohorts of participants, with the first cohort consisting of neurologically and psychiatrically healthy young adults (cohort 1: 117 young), and the other two cohorts including both young and older healthy participants (cohort 2: 58 young, 83 older; cohort 3: 64 young, 84 older). Participants were recruited via billboards, public outreach events of the Leibniz Institute for Neurobiology and local newspapers (for details on recruitment see ^30,57^). All participants were right-handed according to self-report and fluent in German. With the exception of two young participants in cohort 2 (one South Asian, one South American) and one young participant in cohort 3 (Middle Eastern), all participants were of European ancestry. The Mini-International Neuropsychiatric Interview (M.I.N.I.; ^97^; German version by ^98^ was used to exclude present or past psychiatric disorders. Further contraindications for participation included alcohol or drug abuse or the use of neurological or psychiatric medication. Data from sub-cohorts of the current study population have been reported in previous studies (cohort 1: ^99^; cohort 2 and 3:^30,57^). Importantly, the analyses performed here have not been conducted in the previous publications, and cohort 3 included 40 additional participants whose data have not been reported previously. Detailed demographic data are summarized in Table 1.

#### Informed consent and ethics approval

Written informed consent was obtained from all participants in accordance with the Declaration of Helsinki ^100^, and the study was approved by the Ethics Committee of the Otto von Guericke University Magdeburg, Faculty of Medicine (Approval number 33/15).

### Method details

#### Experimental paradigm

During the fMRI experiment, participants performed an incidental visual memory encoding paradigm, using an indoor/outdoor judgment as encoding task ^8^. In cohort 1 ^99,101,102^, the trial timings were slightly shorter compared to cohorts 2 and 3, where we employed the adapted version of the paradigm also used in the DELCODE study ^23,49,103^ (see Table 1 for an overview of differences in trial timings and acquisition parameters). Subjects viewed photographs of indoor and outdoor scenes, which were either novel (44 indoor and outdoor scenes each) or were repetitions of two pre-familiarized “master” images (one indoor and one outdoor scene). Participants performed an “indoor” versus “outdoor” judgment on the images via button press. Each picture was presented for 2.5 s, followed by a variable delay ranging from 0.50 to 2.50 s in cohort 1 and from 0.70 s to 2.65 s in cohorts 2 and 3. The trial timing followed a near-exponential jitter and was optimized to improve estimation of the trial-specific BOLD responses^104^.

Approximately 70 min (cohorts 2, 3) to 90 min (cohort 1) after the start of the fMRI session, participants underwent a computer-based recognition memory test outside the scanner, during which they were presented with photographs from the fMRI encoding phase (*old*; n = 88 plus the two master images), randomly intermixed with previously unseen (*new*) images (n = 44). Participants reported their memory confidence orally on a five-point rating scale from 1 (“definitely new”) to 5 (“definitely old”), and the overt responses were recorded by an experimenter. For details, also see Assmann et al. ^99^ and Soch et al. ^23,30^.

#### MRI data acquisition

Structural and functional MRI data were acquired on two Siemens 3T MR tomographs (cohort 1 and 2: Siemens Verio; cohort 3: Siemens Skyra; see Table 1), using previously reported protocols (cohort 1: see ^99^; cohorts 2 and 3: see ^23^, corresponding to the DELCODE MRI protocol, see ^49,105,106^.

A T1-weighted MPRAGE image (TR = 2.5 s, TE = 4.37 ms, flip-α = 7°; 192 slices, 256 x 256 in-plane resolution, voxel size = 1 x 1 x 1 mm) was acquired for co-registration and improved spatial normalization. In cohorts 2 and 3, phase and magnitude fieldmap images were acquired to improve correction for artifacts resulting from magnetic field inhomogeneities (*unwarping*, see below).

For functional MRI (fMRI), 206 T2*-weighted echo-planar images were acquired in interleaved-ascending slice order (see Table 1 for acquisition details). The complete study protocol also included additional structural MR images, which were not used in the analyses reported here (details available upon request).

#### fMRI data preprocessing

Data preprocessing and analysis was performed using Statistical Parametric Mapping (SPM12; University College London; https://www.fil.ion.ucl.ac.uk/spm/software/spm12/) running on MATLAB R2018b (Mathworks, Natick, MA). EPIs were corrected for acquisition time delay (*slice timing*), head motion (*realignment*) and magnetic field inhomogeneities (*unwarping*), using voxel-displacement maps (VDMs) derived from the fieldmaps (cohorts 2 and 3). The MPRAGE image was spatially co-registered to the mean realigned (cohort 1) or *unwarped* (cohorts 2 and 3) image and *segmented* into six tissue types, using the unified segmentation and normalization algorithm implemented in SPM12. The resulting forward deformation parameters were used to *normalize* EPIs into a standard stereotactic reference frame (Montreal Neurological Institute, MNI; voxel size = 3 x 3 x 3 mm). Normalized images were spatially *smoothed* sing an isotropic Gaussian kernel of 6 mm full width at half maximum (FWHM).

### Additional behavioral measures of memory performance

The Verbal Learning and Memory Test (VLMT) and the Wechsler Memory Scale (WMS), subscale Logical Memory, were administered to the study participants as measures of explicit memory performance independent of the fMRI experiment (for details in cohort 1, see ^101^; for details in cohorts 2 and 3, see ^107^).

#### Verbal Learning and Memory Test (VLMT)

The Verbal Learning and Memory Test (VLMT) consists of two lists of 15 unrelated words, a study list and a distracter list ^37^. The experiment included a learning phase and a recall phase. During the learning phase, participants were shown the words from the first list visually, one after another. When all words had been presented, participants were requested to write down every word they could remember. This procedure was repeated five times consecutively. Next, the distracter list was shown only once, followed by written recall. After presentation and recall of the distracter list, the recall phase began, where participants were asked to write down the words from the initial list. Additional recall phases took place after a 30-minute interval and after 24 hours.

#### Wechsler Memory Scale (WMS), Logical Memory

The Logical Memory subscale of the WMS (Wechsler Memory Scale) was implemented as an auditory version with slight modifications ^38^. Participants were presented with two brief stories via headphones and were instructed to write them down immediately after listening. Recall tests were conducted 30 minutes and 24 hours later. The retrieved stories were evaluated for 25 items, representing details from the stories, by two independent experimenters. Thus, a maximum score of 25 points could be obtained for each story and recall time interval.

### Quantification and statistical analysis

#### General linear model (GLM)-based fMRI data analysis

Statistical analysis of fMRI data was performed based on a two-stage mixed-effects model as implemented in SPM12. At the first stage (single-subject level), we used a parametric general linear model (GLM) of the subsequent memory effect that has previously been demonstrated to outperform the commonly employed categorical models ^23^. The model included two onset regressors, one for novel images (“novelty regressor”) and one for presentations of the two pre-familiarized images (“master regressor”). Both regressors consisted of short box-car stimulus functions (2.5 s), which were convolved with the canonical hemodynamic response function (HRF), as implemented in SPM12.

The regressor reflecting subsequent memory performance was obtained by parametrically modulating the novelty regressor with an arcsine function describing subsequent recognition confidence. Specifically, the parametric modulator (PM) was given by

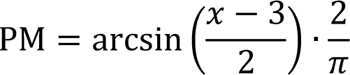

where *x* ∈ {1, 2, 3, 4, 5} is the subsequent memory report, such that – 1 ≤ PM ≤ +1. By employing the arcsine transformation, *definitely* remembered (response “5”) or forgotten (response “1”) items were weighted more strongly than *probably* remembered (response “4”) or forgotten (response “2”) items (^23^, Fig. 2A). The model also included the six rigid-body movement parameters obtained from realignment as covariates of no interest and a constant representing the implicit baseline.

At the second stage of the model (across-subject level), contrasts of interest (novelty contrast: novel vs. master images; memory contrast: parametric memory regressor) were submitted to a one-sample t-test (cohort 1) and to ANCOVA models with age group as between-subjects factor and gender as a binary covariate of no interest (cohort 2, 3), respectively. The significance level was set to p < .05, whole-brain-corrected for family-wise error (FWE) at voxel-level, with a minimum cluster size of k = 10 adjacent voxels.

#### Dynamic Causal Modeling

Effective connectivity analysis was performed using DCM version 12.5 as implemented in SPM12. DCM uses an input-state-output model based on a bilinear state equation:

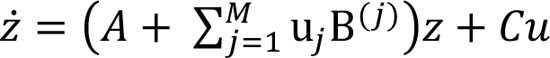

where ͹ is the temporal derivative of the state variable *z*, which describes neuronal activity resulting from intrinsic effective connectivity (*A*), changes in connectivity due to the contextual modulations (*B*) and the direct influence of the driving input *u* (*C*). The thus defined neuronal model is coupled to a biologically plausible neurovascular model of the BOLD response, and the coupled models are used to predict the BOLD time series in *a priori* defined volumes of interest (VOIs).

The goal of our DCM analysis was two-fold:

1. First, we aimed to assess the intrinsic effective connectivity of the PPA, the HC, and the Prc during novelty processing and the connectivity changes related to successful versus unsuccessful encoding of novel visual information.
2. Second, we aimed to assess, which connections within a temporo-parietal memory network are affected by age and whether age-related differences in effective connectivity are associated with memory performance in older adults.

### Definition of regions of interest and time series extraction

ROIs were defined based on both anatomical and functional criteria, following previously described approaches ^32,33^: Anatomical constraints were defined using anatomical ROIs taken from Automated Anatomical Labeling (AAL), a canonical parcellation of the human brain ^108^, as implemented in the WFU PickAtlas (Wake Forest University; https://www.nitrc.org/projects/wfu_pickatlas/) or from previously described literature-based ROIs. Functional constraints were derived of memory and novelty contrasts in a previously published subset of cohorts 2 and 3 (106 young and 111 older participants; ^23,30^). Based on the meta-analytic evidence on encoding-related activations and deactivations ^4^ and the age-related changes of those patterns ^5^, and guided by our own replication of those results, the ROIs were selected as follows:^d^

1. The PPA, which showed a pronounced response during both novelty detection and successful encoding ^30^ was used as the network’s driving input region, given its well-replicated and relatively specific response to scene stimuli ^46^ and particularly to novel scenes ^31,109^. Given that the definition of the PPA is mainly based on its responsivity to scene stimuli rather than on anatomical landmarks, we restricted our ROI to a probabilistic definition of the PPA based on local maxima of activation clusters that were shown to respond to scene stimuli in previous studies (for details see ^110^). This probabilistic ROI was further restricted to the anatomical boundaries of the fusiform and parahippocampal gyri as defined by the AAL parcellation. Considering the intended use of the PPA as input region responding to all presentations of novel scenes (i.e., irrespective of subsequent encoding success), the thresholded SPM for the novelty contrast (novel > master images) was multiplied with the mask of the PPA, yielding a ROI for the PPA.
2. The HC has repeatedly been shown to respond more strongly to subsequently remembered compared to subsequently forgotten information in both young and older adults ^4,5^, a finding replicated in the present study (Figure 2). The ROI for the HC was thus defined by multiplying the thresholded SPM for the memory contrast (positive effects of the parametric memory regressor) with the anatomical boundaries of the HC from the AAL atlas and with a sphere seeded at the local group maximum of the contrast within the right anterior HC ([x y z] = [21-10-19]; r = 15 mm). The focus on the anterior HC was based on previous observations that encoding-related anterior hippocampal activations can be robustly observed in older adults ^5^ and are at the same time sensitive to memory decline in aging ^21^. Furthermore, by restricting the ROI to the anterior HC, we aimed to avoid confounds from potential blurring of activity from the PPA into the posterior hippocampus.
3. The Prc emerged as the region most robustly activating more strongly (or, deactivating less strongly) during successful memory formation in older rather than young adults (Figure 2), in line with previous meta-analytic observations ^5^. We thus inclusively masked the thresholded SPM for the age group contrast (older > young) from the second-level memory GLM with the AAL Prc mask. To exclude activations in more lateral parietal cortex and posterior cingulate cortex, we further restricted the Prc ROI to a sphere centered at the local maximum of the age contrast ([x y z] = [6-64 38]; r = 18 mm). The larger radius compared to the hippocampal ROI was chosen based on the larger size of both the anatomical structure and the activation cluster.

To improve signal-to-noise ratio (SNR), we further restricted the ROIs to voxels with an uncorrected p < 0.25 on the respective contrast in the first-level GLM of each subject. The first eigenvariate time series were extracted from the thus obtained volumes of interest (VOIs) and adjusted for effects of interest (EOI) modeled and explained by the individual first-level GLMs. As agreement of ROI locations across participants and cohorts has been deemed critical for DCM group analyses ^111^, we aimed to ensure that our selection procedure yielded consistent results across cohorts and age groups. To this end, we computed the mean ROI coordinates for each subject and assessed their distribution within and across cohorts and age groups (Supplementary Table S8).

#### First-level DCM analysis

The Parametric Empirical Bayes (PEB) framework implemented in SPM12 ^25,26^ allows for an efficient estimation of effective connectivity at group level by estimating the full model for each individual subject and pruning parameters that do not contribute to the model quality at the second level. In the present study, we employed bilinear DCM as implemented in the DCM PEB framework (^25^; cf. eq. 2}), similarly to previous descriptions of the approach ^112,113^.

The most general model which was estimated for each subject included the ROIs of the HC, PPA, and Prc, assuming full intrinsic connectivity, including self-connections. Novelty (i.e., presentation of novel, but not *master* images) was chosen as driving input to the PPA, and memory (the arcsine-transformed recognition memory responses representing encoding success) was included as a potential contextual modulator at all between-regions connections, but not at the self-connections (Figure 1, right). The reason for not allowing for a modulation of self-connections was that the self-connections themselves turned out to be inconsistent across study cohorts (Figure 4A). Neural responses to the highly familiar *master* images were not included in the model and thus formed the implicit baseline ^25^. The slice timing model implemented in the DCM PEB framework ^114^ was set to the last acquired slice ^25,115^. This was motivated by the fact that the stimulus duration (2.5 s) was close to the TR, and mini-blocks of 2.5 s provided a better model fit than delta functions in first-level GLM analysis ^23^.^e^

Model estimation was performed using variational Laplace ^116^, which provides both, posterior estimates of connection strengths and the free energy approximation to the marginal likelihood^117^. To yield a higher proportion of explained variance at the single-subject level, an iterative fit was performed, using the group mean parameter estimates as empirical priors (SPM function *spm_dcm_peb_fit.m* ^26^).

#### Group-level inference

Group-level inference on the effective connectivity within the temporo-parietal network was performed using Bayesian model reduction (BMR) and averaging (BMA), as implemented in the DCM PEB framework ^26,116^. After estimating each participant’s full model (allowing for all potential connections and contextual modulations), the thus obtained DCMs were submitted to a second-level Bayesian GLM ^26,118^. To assess the strength of effective connectivity between the three ROIs as well as their modulation by successful memory performance and aging, the 15 parameters from model inversion (A-matrix and B-matrix) were submitted to BMR, followed by BMA. BMR compares the full model with a model space of nested models where one or more parameters (i.e., connections or contextual modulations) are switched off, keeping the parameters with the most evidence ^26,116,119^. Subsequently, BMA was employed to average the parameters across models, weighted by the evidence of each model ^26,120^. To test parameters across (or between) groups, we thresholded the posterior probability (PP) at 0.95 for any given parameter (or parameter difference) being larger than zero (*PP* = Pr(θ > 0|y) > 0.95). Parameters with supra-threshold PP in all three cohorts (or in cohorts 2 and 3, when testing for age group effects) were considered robust, and parameters with a PP > 0.95 in two cohorts were considered limited evidence. It must be noted that it is commonly recommended to use a threshold based on free energy, as this approach takes into account the posterior covariance during group model comparison ^26^. However, this might on the downside potentially reduce generalizability due to subtle differences in parameter strength and thus differential pruning of parameters across cohorts. As testing for reproducibility of effects was a main goal of this study, we thus primarily reported marginal posterior probabilities and additionally computed the PPs based on the free energy thresholds for all parameters and age group differences to evaluate the agreement between marginal PPs and the (more stringent) free energy-based threshold.

### Prediction of memory performance from DCM parameters

After group-level PEB analysis, we assessed whether the estimated parameters were associated with memory performance in older adults, and thereby contribute to assess the neurobiological underpinnings of cognitive aging.

#### Memory performance estimate

To obtain a measure of memory performance that takes into account recognition confidence, the area under the curve plotting hits (i.e., correctly recognized previously seen stimuli) against false alarms (i.e., falsely recognized previously unseen stimuli) was calculated as described previously ^30^. To this end, *o*_1_, …, *o*_5_ and *n*_1_, …, *n*_5_ were defined as the numbers of old stimuli and new stimuli, respectively, rated as 1 (“definitely new”) to 5 (“definitely old”) in the delayed recognition test. Hit rates (H) and false alarm (FA) rates as functions of a threshold *t* ∈ {0,1, …, 5} are defined as proportions of old and new stimuli, respectively, rated higher than *t*:

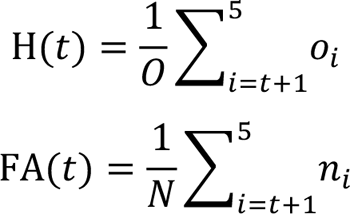

where *O* = *o*_1_ + … +*o*_5_ and *N* = *n*_1_ + … +*n*_5_. Note that H(0) = FA(0) = 1 and H(5) = FA(5) = 0. The hit rate can then be expressed as a function of the FA rate: *y* = *f*(*x*), such that *y* = H(*t*) and *x* = FA(*t*) for each *t* = 0,1, …, 5

The area under the ROC curve (AUC) is then obtained by computing the integral of this function from 0 to 1:

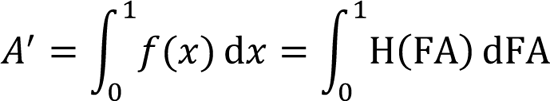

The value of this integral is referred to as AUC or *A*^’^ (“A-prime”) and provides a measure for memory performance, similar to the corrected hit rate, but also accounting for recognition confidence. When responses are completely random, *A*^’^ equals 0.5, reflecting pure guessing. When all old items are recognized and all new items are correctly rejected, *A*^’^ equals 1, reflecting 100% performance. An advantage over the frequently employed *d*’ measure ^121^ is that *A*^’^ has finite values even in cases of zero or perfect performance.

To compare memory performance between young and older adults in cohorts 2 and 3, we computed two-sample t-tests, assuming unequal variances (i.e., Welch’s tests). Bayes factors were computed using the Matlab Bayes Factor Toolbox ^122^.

#### Brain-behavior correlations

Given the high interindividual variability of the DCM parameters, which was suggestive for outliers (Figure 3), we computed outlier-robust Shepherd’s *Pi* correlations ^35^ between connections that showed a robust age-group-related difference – as defined in 2.6.3 – and A’ as a memory measure. Shepherd’s *Pi* correlations have been proposed as a method to improve reliability of brain-behavior correlations, which have been criticized for their lack of robustness ^123^ and susceptibility to outliers ^35,124^. The approach is based on Spearman’s correlation and includes a bootstrap-based estimation of the Mahalanobis distance, thereby allowing for an unbiased detection and exclusion of outliers. To comply with the Bayesian framework of our DCM analyses, Wetzels’ Bayes factors ^36^ were computed for all correlation coefficients, as implemented in the BayesFactor Toolbox for MATLAB ^122^ (https://github.com/klabhub/bayesFactor). As the sample size is reduced by the outlier exclusion of the Shepherd’s *Pi* correlation, all Bayes factors were computed based on the effective sample size (i.e., not counting outliers). First, a likelihood function *L* was defined as described previously (see Wetzels & Wagenmakers^36^, eq 13):

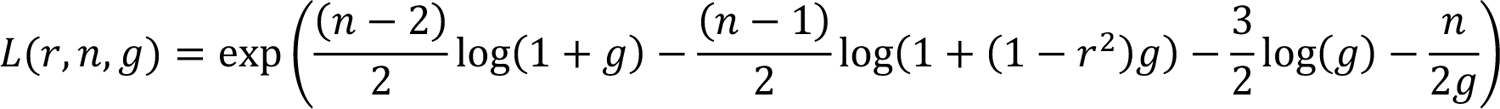

where n is the sample size, r is the correlation coefficient, and *g* represents Zellner’s g prior^125,126^.

The Bayes factor BF_10_, quantifying the relative likelihood of the alternative hypothesis over the null hypothesis, was computed by integrating the likelihood function over *g* and multiplying with a sample-size-dependent scaling factor:

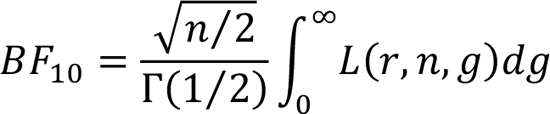

where Γ(*x*) is the gamma function.

To further explore which DCM parameters were associated with memory performance in older and also young adults, we computed Bayesian Shepherd’s *Pi* correlations between all DCM parameters and A’ as the memory measure of interest as well as the performance measures from two independent memory tests (VLMT, WMS; see 7.4.4 for details).

We then estimated the overall evidence for the correlations across the study cohorts by averaging the correlation coefficients weighted by their effective sample sizes across cohorts, separately for young and older adults. To this end, correlation coefficients were first submitted to Fisher’s z transformation to make them approximately normally distributed. Next, we computed the weighted mean z values

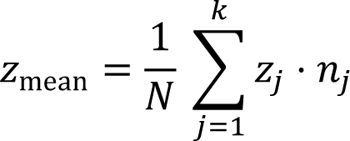

where *Z*_mean_ is the weighted mean z-transformed correlation coefficient, N is the total effective sample size across all cohorts, k is the number of cohorts, z_j_ is the j^th^ z-transformed correlation coefficient, and n_j_ is the j^th^ effective sample size (i.e., the sample size after outlier exclusion by the Shepherd’s *Pi* correlation). Weighted mean z-values were then transformed to the corresponding correlation coefficients using the inverse Fisher’s z-transformation. Next, we computed the combined Bayes factors from all samples as described above, using N (i.e., the total effective sample size over all cohorts) as sample size (for a similar approach with t-test statistics, see Nikolakopoulos & Ntzoufras^83^). The resulting weighted mean correlation coefficients and cumulative Bayes factors are reported in Supplementary Tables S9 and S10.

## 9. Declaration of Interest

The authors declare no competing interests.

## Footnotes

a In cohort 2, a single voxel in the rostral anterior cingulate showed a gender difference in the subsequent memory contrast, with women showing lower encoding-related deactivations compared to men.

b The effective sample size was defined as the sample size remaining after exclusion of i) missing data points from individual participants (e.g., due to skipping of the 24 h delayed recall) and ii) the Bootstrap-based outlier exclusion implemented in the Shepherd’s Pi correlation.

c In the combined analyses across cohorts, there was some evidence for an association between the driving input to the PPA during novelty processing and memory performance in the independent tests in older adults. Please see the Discussion for a more detailed discussion of this observation.

d Previous DCM studies of memory encoding have also included prefrontal regions in their models. In the course of the present study, we also tested models that the medial prefrontal cortex (mPFC), however, this did not improve the model fit. Please see also the Discussion section.

e As a control, we also computed DCMs using TR/2 and TR/_*N*slices_ (i.e., number of slices) in the slice timing model. Explained variance of the first-level DCM analyses was substantially poorer for TR/_*N*slices_ (cohort 1: p = .001; cohort 2: p = .057; cohort 3: p < .001) and did not differ significantly for TR/2 (all > .161). Results of second- level PEB analysis for the model using TR/2 were qualitatively highly similar to the results of that using TR reported here.

## Notes

### Competing Interest Statement

The authors have declared no competing interest.

### Summary of Updates

Fully Bayesian statistical approach has been implemented for the brain-behavior correlations.

