## supplementary figures and tables for "Inhibitory temporo-parietal effective connectivity is associated with explicit memory performance in older adults"

### - Supporting Online Material –

#### 1. Supplementary Figures

**Figure S1.** *Illustration of the mean centers of the ROIs in relation to the corresponding (functional) anatomical structures.* (related to Figure 1, Figure 6 and Results section "Location of regions of interest")

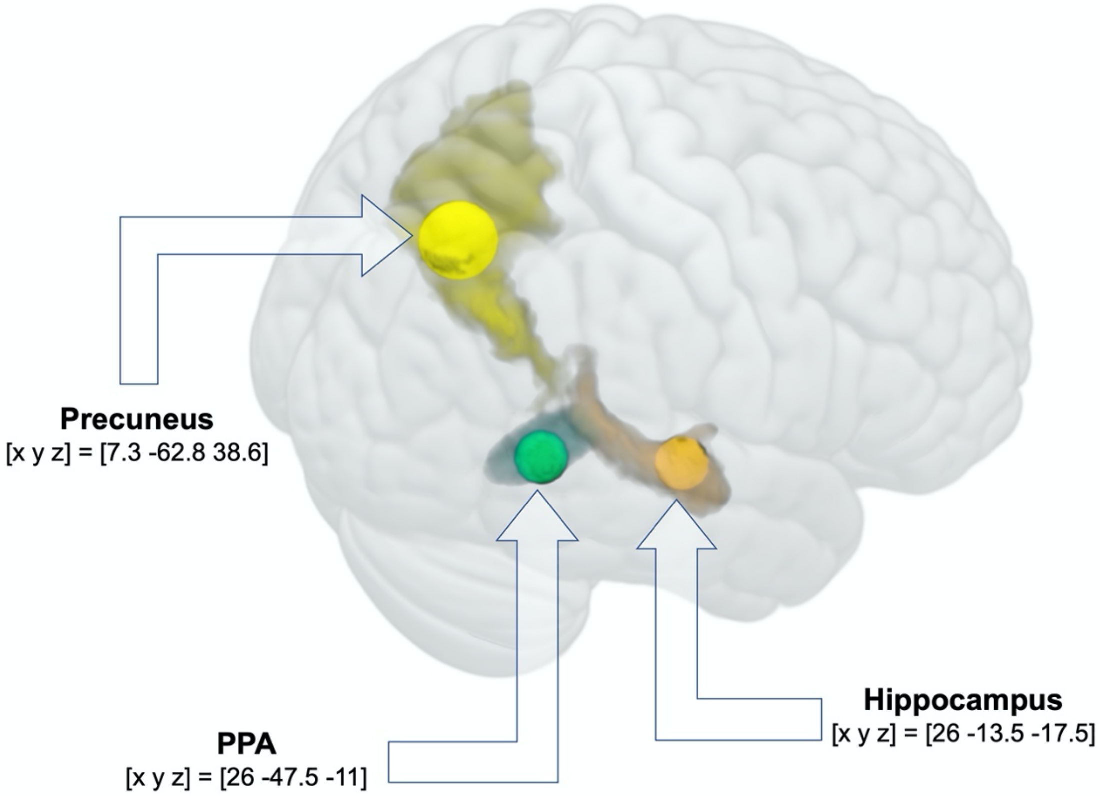


Anatomical structures of the hippocampus and precuneus were generated using Automated Anatomical Labeling (AAL), as implemented in the WFU Pickatlas. The ellipsoid functional anatomical ROI of the PPA was based on coordinates obtained from independent studies and has been described previously (Zweynert et al., 2011).

**Figure S2.** *Replicability of DCM results across cohorts.* (related to Figure 3, Figure 4 and Table 2)

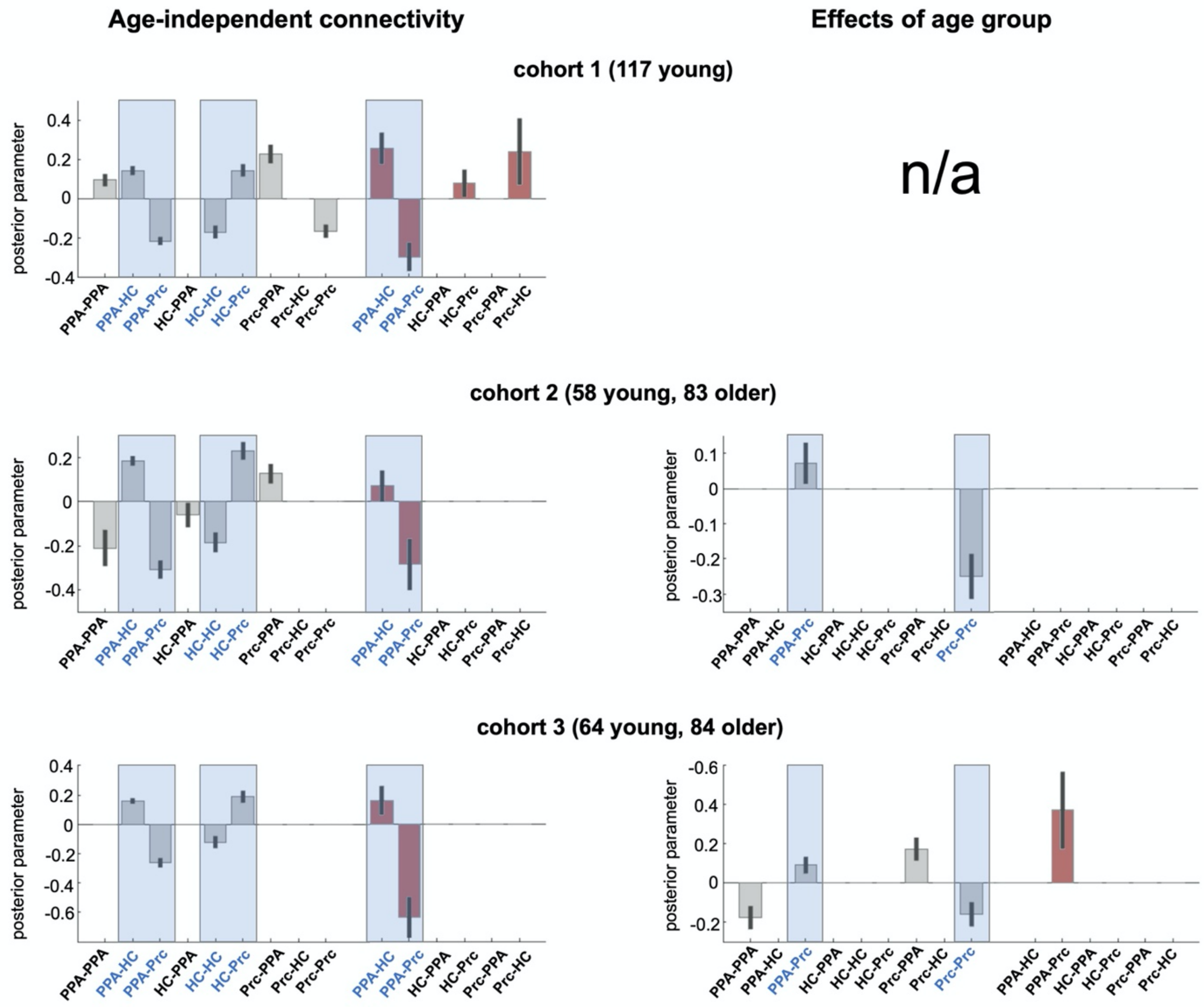


Bar plots show group-level Bayesian mean parameter estimates and 90% confidence intervals. Grey bars denote intrinsic connections (A-parameters) and red bars denote contextual modulations (B-parameters). Plots highlighted in blue show connections that were replicated across all three cohorts (main effects, left panel) and in cohorts 2 and 3 (age group effects, i.e., older - young, right panel), respectively.

**Figure S3.** *Correlations of DCM parameters and memory performance in young and older participants.* (related to Figure 5)

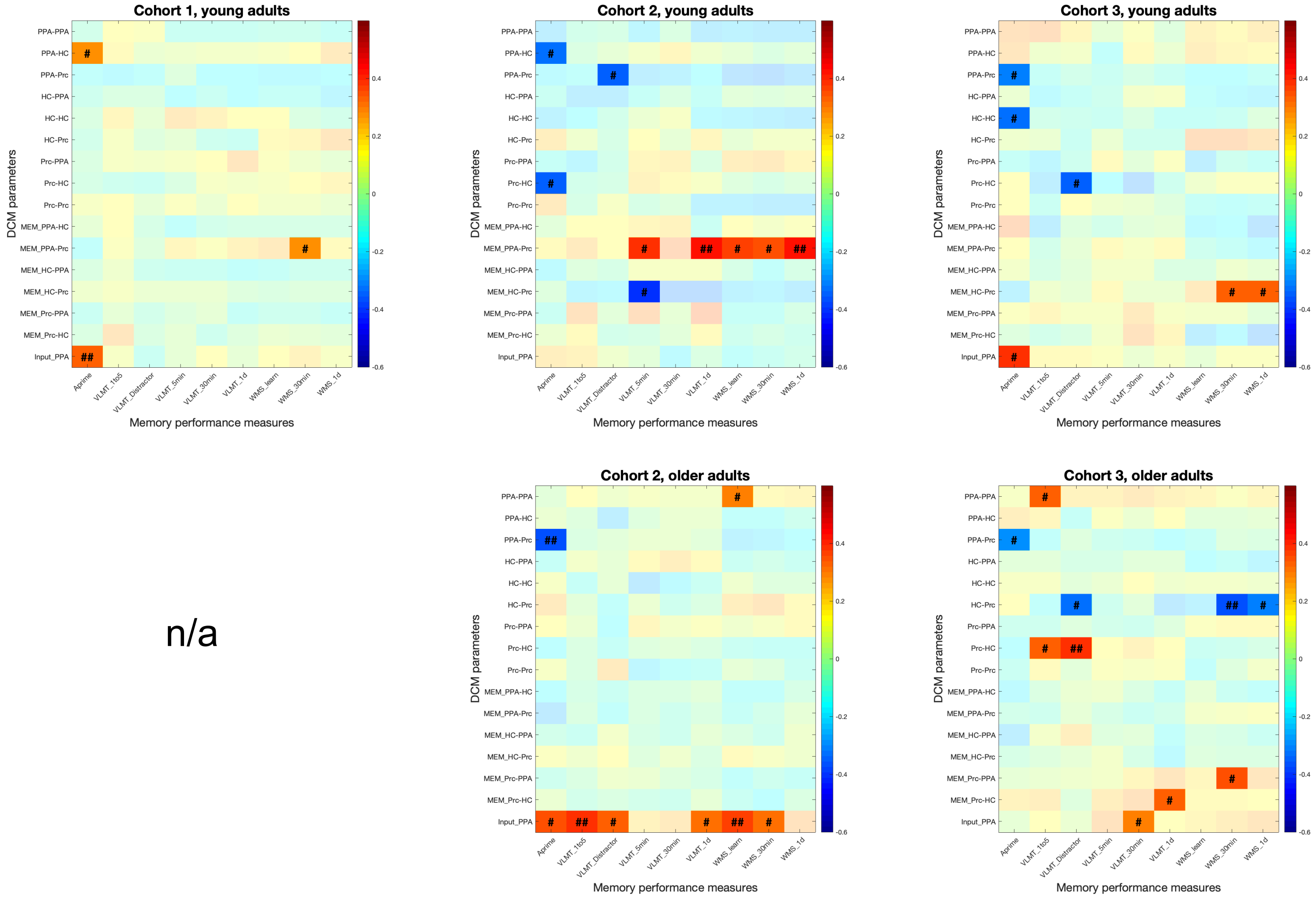


The figure displays the heatmap of Bayesian Shepherd's *Pi* correlations between DCM parameters (rows) and behavioral measures of memory (A', VLMT, WMS), separated by age cohorts and age groups. Correlations with Bayes factor > 2 are highlighted. $^{\#}$ $BF_{10} > 2$; $^{\#\#}$ $BF_{10} > 10$.

#### 2. Supplementary Tables

**Table S1.** *Memory performance, separated by gender (young and older adults).*
(related to Table 1)

| | **Cohort 1** | **Cohort 2** | **Cohort 3** |
|---|---|---|---|
| **A' young** | | | |
| ***female*** | 0.81 +/- 0.06 | 0.84 +/- 0.07 | 0.83 +/- 0.07 |
| ***male*** | 0.80 +/- 0.09 | 0.82 +/- 0.07 | 0.80 +/- 0.07 |
| **T** | 0.93 | 1.16 | 1.72 |
| **$BF_{10}$** | 0.291 | 0.466 | 0.908 |
| **A' older** | | | |
| ***female*** | n/a | 0.79 +/- 0.06 | 0.76 +/- 0.08 |
| ***male*** | n/a | 0.76 +/- 0.09 | 0.78 +/- 0.06 |
| **T** | n/a | 1.47 | -1.11 |
| **$BF_{10}$** | n/a | 0.591 | 0.402 |

Data are shown separately for each cohort and age group. A' = area under the curve in a ROC analysis of the recognition memory test, reflecting behavioral accuracy (hits vs. false alarms). Please note that gender is denoted as female and male only, as no participants identified as other.

**Table S2.** *Effects of novelty (novel vs. master images) in cohort 1 (young adults only).* (related to Figure 2)

| | Hemisphere | Cluster size | peak *t* | peak *p* | x y z |
|---|---|---|---|---|---|
| **Positive effect** | | | | | |
| Lingual gyrus | R | 8369 | 24.00 | <.001 | 27 -46 -10 |
| Fusiform gyrus | L | | 23.04 | <.001 | -27 -55 -13 |
| Lingual gyrus | R | | 20.65 | <.001 | 24 -61 -10 |
| Precentral gyrus | L | 150 | 10.23 | <.001 | -36 5 29 |
| | L | | 10.02 | <.001 | -45 8 29 |
| Mid cingulate gyrus | R | 41 | 8.22 | <.001 | 9 -40 50 |
| Inferior frontal gyrus, pars orbitalis | L | 31 | 8.21 | <.001 | -33 35 -13 |
| Inferior frontal gyrus, pars triangularis | L | 50 | 8.01 | <.001 | -51 35 14 |
| Frontal operculum | R | 67 | 7.96 | <.001 | 39 8 29 |
| Supplementary motor cortex | L | 163 | 7.75 | <.001 | 0 14 50 |
| Mid cingulate gyrus | R | | 7.45 | <.001 | 9 17 44 |
| Mid cingulate gyrus | R | | 5.60 | .004 | 3 23 35 |
| Precentral gyrus | L | 137 | 7.51 | <.001 | -39 -19 56 |
| Postcentral gyrus | L | | 6.61 | <.001 | -45 -28 50 |
| | L | | 6.58 | <.001 | -36 -34 59 |
| Cerebellum | L | 10 | 7.43 | <.001 | -18 -40 -46 |
| Frontal operculum | R | 17 | 7.03 | <.001 | 30 35 -13 |
| Superior frontal gyrus | L | 64 | 6.90 | <.001 | -21 5 56 |
| | L | | 5.70 | .003 | -21 5 47 |
| Cerebellum / vermis | L/R | 13 | 6.82 | <.001 | 0 -52 -37 |
| Mid cingulate gyrus | L | 20 | 6.70 | <.001 | -12 -34 47 |
| Insula | L | 13 | 6.18 | <.001 | -30 26 2 |
| Middle frontal gyrus | R | 39 | 6.15 | <.001 | 24 -1 50 |
| Superior frontal gyrus | R | | 5.95 | .001 | 24 8 53 |
| | R | | 5.29 | .015 | 27 -1 59 |
| Inferior frontal gyrus, pars triangularis | R | 13 | 5.71 | .002 | 45 32 11 |
| **Negative effect** | | | | | |
| Precuneus | R | 780 | 18.25 | <.001 | 6 -64 35 |
| Mid cingulate gyrus | R | | 10.47 | <.001 | 3 -43 23 |
| Posterior cingulate gyrus | R | | 8.44 | <.001 | 3 -28 29 |
| Angular gyrus | L | 485 | 12.78 | <.001 | -45 -67 44 |
| | L | | 10.59 | <.001 | -39 -58 35 |
| Middle temporal gyrus | L | | 7.63 | <.001 | -54 -58 17 |
| Angular gyrus | R | 837 | 12.54 | <.001 | 45 -64 44 |
| | R | | 10.84 | <.001 | 60 -55 23 |
| Middle temporal gyrus | R | | 9.92 | <.001 | 69 -37 2 |
| Posterior cingulate gyrus | L | 46 | 9.55 | <.001 | -18 -43 11 |
| Precuneus | R | 52 | 9.24 | <.001 | 21 -40 14 |
| Thalamus | R | | 5.94 | .001 | 9 -31 11 |
| Middle frontal gyrus | L | 47 | 7.86 | <.001 | -36 20 47 |
| | L | | 6.04 | .001 | -42 23 38 |
| Superior frontal gyrus | R | 39 | 7.26 | <.001 | 12 41 44 |
| Middle frontal gyrus | L | 12 | 6.12 | <.001 | -36 53 2 |
| Middle temporal gyrus | L | 40 | 6.09 | <.001 | -66 -37 -1 |
| | L | | 5.91 | .001 | -66 -40 8 |
| Superior temporal gyrus | L | | 5.33 | .013 | -63 -52 11 |
| Middle frontal gyrus | R | 36 | 6.07 | <.001 | 36 17 53 |
| | R | | 5.97 | .001 | 39 14 44 |
| Superior frontal gyrus | L | 13 | 5.99 | .001 | -18 59 2 |

Coordinates are given in MNI space, normalized voxel size = 3x3x3 mm. All p-values are corrected for family-wise error (FWE) at peak level, minimum cluster size = 10 voxels.

**Table S3.** *Effects of novelty (novel vs. master images) in cohort 2 (young and older adults).* (related to Figure 2)

| | Hemisphere | Cluster size | peak *t* | peak *p* | x y z |
|---|---|---|---|---|---|
| **Young adults** | | | | | |
| Lingual gyrus | L | 6047 | 19.29 | <.001 | -27 -52 -7 |
| Fusiform gyrus | R | | 19.07 | <.001 | 30 -52 -7 |
| Lingual gyrus | L | | 17.49 | <.001 | -24 -70 -10 |
| Cerebellum | L | 68 | 10.79 | <.001 | -15 -46 -46 |
| Cerebellum | R | 56 | 9.18 | <.001 | 18 -40 -46 |
| Cerebellum / vermis | L/R | 33 | 8.31 | <.001 | 0 -55 -37 |
| Frontal operculum | L | 89 | 6.90 | <.001 | -42 8 26 |
| Precentral gyrus | L | | 6.22 | <.001 | -42 2 38 |
| Supplementary motor cortex | R | 36 | 6.60 | <.001 | 3 14 47 |
| Frontal operculum | R | 36 | 6.44 | <.001 | 42 8 26 |
| Orbitofrontal cortex | L | 11 | 5.92 | .001 | -36 32 -16 |
| **Older adults** | | | | | |
| Fusiform gyrus | L | 8072 | 20.69 | <.001 | -27 -46 -10 |
| Lingual gyrus | L | | 20.54 | <.001 | -27 -82 23 |
| Fusiform / parahippocampal gyrus | R | | 20.18 | <.001 | 27 -40 -13 |
| Cerebellum | L | 87 | 12.91 | <.001 | -15 -43 -46 |
| Precentral gyrus | L | 509 | 10.77 | <.001 | -42 5 29 |
| Superior frontal gyrus | L | | 8.94 | <.001 | -21 2 53 |
| Precentral gyrus | L | | 8.69 | <.001 | -36 -13 47 |
| Cerebellum | R | 54 | 9.48 | <.001 | 15 -43 -49 |
| Supplementary motor cortex | R | 289 | 9.27 | <.001 | 6 17 47 |
| | L | | 8.19 | <.001 | -6 8 50 |
| Mid cingulate cortex | R | | 6.70 | <.001 | 12 23 29 |
| Inferior frontal gyrus, pars orbitalis | R | 93 | 9.25 | <.001 | 30 32 -10 |
| Insula | R | | 7.24 | <.001 | 33 26 2 |
| | R | | 5.72 | .002 | 39 17 2 |
| Middle frontal gyrus | R | 168 | 8.81 | <.001 | 24 -1 50 |
| Precentral gyrus | R | | 8.62 | <.001 | 39 -10 50 |
| Inferior frontal gyrus, pars orbitalis | L | 131 | 8.60 | <.001 | -30 32 -13 |
| Insula | L | | 7.38 | <.001 | -30 26 -1 |
| Frontal operculum | R | 136 | 8.37 | <.001 | 42 11 29 |
| Inferior frontal gyrus, pars triangularis | R | 52 | 6.77 | <.001 | 48 35 14 |
| Cerebellum | L | 12 | 6.64 | <.001 | -27 -67 -49 |
| Inferior frontal gyrus, pars triangularis | L | 38 | 6.39 | <.001 | -48 35 11 |
| | L | | 5.84 | .001 | -51 32 20 |
| **Young > older adults** | | | | | |
| Lingual / fusiform gyrus | L | 26 | 7.01 | <.001 | -24 -52 -4 |
| | L | | 5.54 | .004 | -18 -52 -10 |
| Fusiform / parahippocampal gyrus | R | 33 | 6.79 | <.001 | 30 -49 -4 |

Coordinates are given in MNI space, normalized voxel size = 3x3x3 mm. All p-values are corrected for family-wise error (FWE) at peak level, minimum cluster size = 10 voxels.

**Table S4.** *Effects of novelty (novel vs. master images) in cohort 3 (young and older adults).* (related to Figure 2)

| | Hemisphere | Cluster size | peak *t* | peak *p* | x y z |
|---|---|---|---|---|---|
| **Young adults** | | | | | |
| Lingual gyrus | R | 7482 | 19.21 | <.001 | 30 -46 -7 |
| Fusiform / parahippocampal gyrus | R | | 19.10 | <.001 | 30 -40 -13 |
| Lingual gyrus | R | | 18.97 | <.001 | 27 -55 -10 |
| Cerebellum | R | 52 | 8.98 | <.001 | 15 -46 -49 |
| | R | | 8.16 | <.001 | 21 -40 -46 |
| Supplementary motor cortex | L | 103 | 8.06 | <.001 | -3 11 50 |
| Medial frontal gyrus | R | | 6.18 | <.001 | 3 20 41 |
| Precentral gyrus | L | 90 | 7.69 | <.001 | -45 5 29 |
| Frontal operculum | L | | 7.13 | <.001 | -36 5 26 |
| Middle frontal gyrus | R | 42 | 7.64 | <.001 | 27 2 50 |
| Superior frontal gyrus | L | 65 | 7.64 | <.001 | -24 2 53 |
| Frontal operculum | R | 30 | 7.43 | <.001 | 42 8 26 |
| Precentral gyrus | L | 111 | 7.39 | <.001 | -33 -22 56 |
| | L | | 6.33 | <.001 | -42 -28 53 |
| | L | | 5.53 | .004 | -33 -34 44 |
| Inferior frontal gyrus, pars triangularis | L | 32 | 7.10 | <.001 | -51 32 11 |
| Inferior frontal gyrus, pars orbitalis | R | 37 | 6.85 | <.001 | 30 32 -13 |
| Insula | L | 68 | 6.32 | <.001 | -30 17 2 |
| Inferior frontal gyrus, pars orbitalis | L | | 5.77 | .001 | -30 26 -13 |
| | L | | 5.46 | .005 | -39 35 -13 |
| **Older adults** | | | | | |
| Middle occipital gyrus | R | 8862 | 20.51 | <.001 | 39 -82 20 |
| Fusiform gyrus | R | | 19.65 | <.001 | 30 -49 -13 |
| Fusiform / parahippocampal gyrus | R | | 19.57 | <.001 | 30 -40 -13 |
| Frontal operculum | R | 260 | 10.53 | <.001 | 42 8 29 |
| Inferior frontal gyrus, pars triangularis | R | | 7.71 | <.001 | 48 32 17 |
| Frontal operculum | L | 461 | 10.53 | <.001 | -39 8 26 |
| Inferior frontal gyrus, pars triangularis | L | | 8.81 | <.001 | -42 29 17 |
| Precentral / postcentral gyrus | L | | 8.20 | <.001 | -42 -10 47 |
| Supplementary motor cortex | R | 246 | 9.63 | <.001 | 6 5 53 |
| | L | | 8.54 | <.001 | -3 11 50 |
| | R | | 8.39 | <.001 | 6 14 47 |
| Inferior frontal gyrus, pars orbitalis | L | 122 | 8.12 | <.001 | -30 29 -7 |
| Orbitofrontal cortex | L | | 7.15 | <.001 | -33 32 -16 |
| Insula | L | | 6.88 | <.001 | -30 17 -1 |
| Insula | R | 87 | 7.62 | <.001 | 33 23 2 |
| Inferior frontal gyrus, pars orbitalis | R | | 7.09 | <.001 | 33 32 -10 |
| Superior frontal gyrus | R | 128 | 7.49 | <.001 | 24 2 50 |
| Precentral gyrus | R | | 6.87 | <.001 | 42 -10 47 |
| Superior frontal gyrus | R | | 6.76 | <.001 | 33 -1 59 |
| **Young > older adults** | | | | | |
| | no voxels survive statistical threshold p < .05, whole-brain FWE-corrected | | | | |

Coordinates are given in MNI space, normalized voxel size = 3x3x3 mm. All p-values are corrected for family-wise error (FWE) at peak level, minimum cluster size = 10 voxels.

**Table S5.** *Effects of successful encoding (parametric) in cohort 1 (young adults only).* (related to Figure 2)

| | Hemisphere | Cluster size | peak t | peak p | x y z |
|---|---|---|---|---|---|
| **Positive effect** | | | | | |
| Middle occipital gyrus | R | 5989 | 16.97 | <.001 | 30 -76 29 |
| | R | | 16.06 | <.001 | 42 -79 23 |
| Middle temporal gyrus | R | | 15.43 | <.001 | 48 -73 20 |
| Frontal operculum | R | 304 | 11.70 | <.001 | 42 8 26 |
| Inferior frontal gyrus, pars triangularis | R | | 11.19 | <.001 | 48 32 17 |
| | | | 8.61 | <.001 | 54 38 11 |
| Inferior frontal gyrus, pars triangularis | L | 262 | 10.73 | <.001 | -42 11 26 |
| | | | 7.45 | <.001 | -42 26 20 |
| Inferior frontal gyrus, pars orbitalis | L | 50 | 10.10 | <.001 | -33 35 -13 |
| Inferior frontal gyrus, pars orbitalis | R | 44 | 9.12 | <.001 | 33 35 -13 |
| Hippocampus | R | 33 | 7.54 | <.001 | 18 -7 -16 |
| Parahippocampal gyrus | R | | 6.09 | <.001 | 21 -19 -19 |
| Rectus gyrus | L | 76 | 7.08 | <.001 | -3 41 -16 |
| Rectus gyrus | R | | 6.91 | <.001 | 6 41 -16 |
| Inferior frontal gyrus, pars orbitalis | L | | 6.39 | <.001 | -9 35 -13 |
| Middle temporal gyrus | R | 15 | 6.50 | <.001 | 51 -7 -19 |
| **Negative effect** | | | | | |
| Inferior parietal gyrus | R | 576 | 13.18 | <.001 | 54 -49 41 |
| Supramarginal gyrus | R | | 11.78 | <.001 | 54 -40 41 |
| Precuneus | L | 865 | 12.56 | <.001 | 0 -73 44 |
| Mid cingulate gyrus | R | | 12.50 | <.001 | 3 -22 38 |
| Precuneus | R | | 12.26 | <.001 | 9 -67 35 |
| Middle frontal gyrus | R | 814 | 10.90 | <.001 | 39 26 41 |
| | R | | 9.64 | <.001 | 39 56 5 |
| | R | | 9.22 | <.001 | 39 35 32 |
| Inferior parietal gyrus | L | 446 | 10.28 | <.001 | -48 -58 41 |
| Inferior parietal gyrus | L | | 9.97 | <.001 | -54 -49 41 |
| Supramarginal gyrus | L | | 8.96 | <.001 | -63 -40 32 |
| Anterior cingulate | R | 391 | 9.63 | <.001 | 3 38 17 |
| | R | | 8.60 | <.001 | 6 44 2 |
| Medial frontal gyrus | R | | 7.86 | <.001 | 3 32 32 |
| Middle frontal gyrus | L | 510 | 9.58 | <.001 | -36 50 11 |
| | | | 8.15 | <.001 | -33 26 38 |
| | | | 8.10 | <.001 | -36 38 29 |
| Superior temporal gyrus | R | 119 | 8.92 | <.001 | 63 -22 -7 |
| Middle temporal gyrus | R | | 6.97 | <.001 | 54 -25 -7 |
| Superior frontal gyrus | | 125 | 8.44 | <.001 | 18 17 59 |
| Medial frontal gyrus | | | 6.51 | <.001 | 9 35 56 |
| Posterior cingulate | R | 11 | 6.80 | <.001 | 18 -40 14 |
| Inferior frontal gyrus, pars triangularis | R | 20 | 5.82 | .001 | 51 20 2 |
| Frontal operculum | R | 10 | 5.70 | .002 | 54 11 8 |

Coordinates are given in MNI space, normalized voxel size = 3x3x3 mm. All p-values are corrected for family-wise error (FWE) at peak level, minimum cluster size = 10 voxels.

**Table S6.** *Effects of successful encoding (parametric) in cohort 2 (young and older adults).* (related to Figure 2)

| | **Hemisphere** | **Cluster size** | **peak *t*** | **peak *p*** | **x y z** |
|---|---|---|---|---|---|
| **Young adults** | | | | | |
| Fusiform gyrus | L | 4348 | 14.15 | <.001 | -27 -52 -10 |
| Fusiform / parahippocampal gyrus | R | | 13.02 | <.001 | 30 -40 -13 |
| Fusuform / parahippocampal gyrus | L | | 12.31 | <.001 | -27 -40 -13 |
| Frontal operculum | R | 116 | 9.27 | <.001 | 45 8 26 |
| Inferior frontal gyrus, pars triangularis | R | 78 | 8.55 | <.001 | 45 32 14 |
| Frontal operculum | L | 75 | 8.01 | <.001 | -39 8 26 |
| Inferior frontal gyrus, pars triangularis | L | 47 | 6.77 | <.001 | -45 29 14 |
| Cerebellum | L | 23 | 6.60 | <.001 | -18 -43 -46 |
| Orbitofrontal cortex | L | 13 | 6.49 | <.001 | -36 35 -16 |
| Inferior frontal gyrus, pars orbitalis | L | | 5.21 | .017 | -27 29 -13 |
| Cuneus | R | 12 | 5.53 | .004 | 12 -91 17 |
| **Older adults** | | | | | |
| Middle occipital gyrus | R | 1269 | 10.51 | <.001 | 42 -76 23 |
| Superior occipital gyrus | R | | 9.74 | <.001 | 24 -64 47 |
| Parahippocampal gyrus | R | | 9.46 | <.001 | 30 -37 -13 |
| Middle occipital gyrus | L | 1132 | 10.35 | <.001 | -36 -82 29 |
| Inferior temporal gyrus | L | | 9.69 | <.001 | -45 -58 -10 |
| Fusiform / parahippocampal gyrus | L | | 9.38 | <.001 | -27 -34 -19 |
| Precuneus | L | 66 | 8.87 | <.001 | -6 -55 11 |
| Precuneus | R | 90 | 8.14 | <.001 | 12 -52 11 |
| Medial occipital cortex | R | | 7.17 | <.001 | 21 -55 17 |
| Precentral gyrus | L | 61 | 7.71 | <.001 | -39 5 29 |
| Inferior frontal gyrus, pars orbitalis | R | 30 | 7.38 | <.001 | 30 32 -13 |
| Orbitofrotal cortex | L | 21 | 7.05 | <.001 | -36 32 -16 |
| Rectus_L | L | 16 | 7.04 | <.001 | -3 35 -16 |
| Frontal operculum | R | 13 | 6.56 | <.001 | 42 5 26 |
| Hippocampus | L | 12 | 6.01 | <.001 | -21 -10 -19 |
| Superior frontal gyrus | L | 11 | 5.74 | .001 | -24 14 44 |
| **Young > older adults** | | | | | |
| Fusiform gyrus | L | 194 | 9.26 | <.001 | -27 -55 -10 |
| | L | | 6.60 | <.001 | -24 -70 -7 |
| Lingual gyrus | R | 181 | 7.10 | <.001 | 30 -46 -7 |
| Fusiform / parahippocampal gyrus | R | | 6.74 | <.001 | 33 -40 -13 |
| Lingual gyrus | R | | 6.71 | <.001 | 27 -61 -7 |
| Middle occipital gyrus | R | 78 | 6.76 | <.001 | 33 -76 38 |
| | R | | 6.08 | <.001 | 30 -73 26 |
| Medial occipital cortex | R | 20 | 6.08 | <.001 | 18 -49 8 |
| Middle occipital gyrus | L | 10 | 5.92 | .001 | -33 -79 23 |
| Frontal operculum | R | 16 | 5.87 | .001 | 48 11 26 |
| Inferior temporal gyrus | R | 23 | 5.68 | .002 | 48 -61 -7 |
| Inferior occipital gyrus | R | | 5.65 | .002 | 39 -64 -13 |
| **Young < older adults** | | | | | |
| Precuneus | R | 58 | 6.43 | <.001 | 6 -64 41 |

Coordinates are given in MNI space, normalized voxel size = 3x3x3 mm. All p-values are corrected for family-wise error (FWE) at peak level, minimum cluster size = 10 voxels.

**Table S7.** *Effects of successful encoding (parametric) in cohort 3 (young and older adults).* (related to Figure 2)

| | Hemisphere | Cluster size | peak *t* | peak *p* | x y z |
|---|---|---|---|---|---|
| **Young adults** | | | | | |
| Middle occipital gyrus | R | 2196 | 14.04 | <.001 | 30 -76 32 |
| Parahippocampal gyrus | R | | 12.15 | <.001 | 30 -37 -13 |
| Inferior temporal gyrus | R | | 11.65 | <.001 | 45 -58 -7 |
| Fusiform gyrus | L | 1946 | 13.56 | <.001 | -30 -40 -13 |
| Middle occipital gyrus | L | | 13.14 | <.001 | -27 -76 32 |
| Inferior occipital gyrus | L | | 11.89 | <.001 | -42 -70 -7 |
| Frontal operculum | R | 210 | 9.90 | <.001 | 39 8 26 |
| Inferior frontal gyrus, pars triangularis | R | | 8.77 | <.001 | 48 32 17 |
| Medial occipital cortex | L | 112 | 9.38 | <.001 | -15 -55 11 |
| Hippocampus | R | 32 | 7.44 | <.001 | 21 -10 -19 |
| Precentral gyrus | L | 38 | 6.96 | <.001 | -42 5 29 |
| Inferior frontal gyrus, pars triangularis | L | | 5.32 | .009 | -45 17 23 |
| Inferior frontal gyrus, pars orbitalis | R | 15 | 6.42 | <.001 | 33 35 -13 |
| Cerebellum | L | 12 | 6.39 | <.001 | -18 -40 -46 |
| **Older adults** | | | | | |
| Middle occipital gyrus | R | 1081 | 10.49 | <.001 | 36 -85 17 |
| | R | | 9.73 | <.001 | 30 -76 32 |
| Fusiform gyrus | R | | 8.58 | <.001 | 27 -46 -13 |
| Middle occipital gyrus | L | 1041 | 10.13 | <.001 | -27 -70 32 |
| | L | | 9.49 | <.001 | -30 -82 29 |
| | L | | 9.34 | <.001 | -42 -79 26 |
| Precuneus | L | 87 | 8.69 | <.001 | -9 -55 14 |
| Precuneus | R | 69 | 7.19 | <.001 | 12 -52 11 |
| Frontal operculum | L | 27 | 6.57 | <.001 | -36 8 26 |
| **Young > older adults** | | | | | |
| Fusiform / parahippocampal gyrus | L | 111 | 6.58 | <.001 | -30 -40 -13 |
| Fusiform gyrus | L | | 6.41 | <.001 | -33 -64 -10 |
| Lingual / fusiform gyrus | L | | 5.79 | .001 | -27 -52 -7 |
| Superior occipital gyrus | R | 29 | 5.86 | .001 | 27 -73 32 |
| Middle occipital gyrus | R | | 5.75 | .001 | 30 -82 32 |
| Inferior temporal gyrus | R | 11 | 5.79 | .001 | 45 -58 -7 |
| Parahippocampal gyrus | R | 13 | 5.69 | .002 | 27 -37 -13 |
| | R | | 5.19 | .017 | 30 -43 -7 |
| Middle occipital gyrus | L | 15 | 5.53 | .003 | -24 -73 26 |
| **Young < older adults** | | | | | |
| Precuneus | R | 312 | 8.75 | <.001 | 3 -64 38 |
| Posterior cingulate gyrus | R | | 6.44 | <.001 | 3 -43 23 |
| | R | | 6.18 | <.001 | 9 -49 29 |
| Mid cingulate gyrus | L | 45 | 6.77 | <.001 | -3 -19 29 |
| Angular gyrus | R | 24 | 6.40 | <.001 | 51 -52 29 |
| Rostral anterior cingulate gyrus | L/R | 23 | 5.96 | <.001 | 0 44 5 |
| Medial frontal gyrus | L/R | | 5.12 | .023 | 0 53 5 |

Coordinates are given in MNI space, normalized voxel size = 3x3x3 mm. All p-values are corrected for family-wise error (FWE) at peak level, minimum cluster size = 10 voxels.

**Table S8.** *Location and size of the regions of interest (ROIs).*
(related to Figure 1; Results section "Locations of regions of interest")

| | Cohort 1 | Cohort 2 | Cohort 3 |
|---|---|---|---|
| **PPA** | | | |
| ***young*** | | | |
| mean [x y z] | **[26.0 -47.6 -11.2]** | **[25.9 -47.3 -11.0]** | **[25.9 -47.3 -11.0]** |
| ± SD | ± [0.29 0.87 0.50] | ± [0.17 0.42 0.33] | ± [0.21 0.50 0.34] |
| size (voxels) ± SD | **54.6** (± 7.48) | **59.3** (± 3.78) | **58.7** (± 5.24) |
| ***older*** | | | |
| mean [x y z] | – | **[26.0 -47.5 -11.2]** | **[25.9 -47.5 -11.1]** |
| ± SD | | ± [0.24 0.68 0.42] | ± [0.20 0.57 0.37] |
| size (voxels) ± SD | – | **56.5** (±5.51) | **56.8** (± 5.57) |
| **Hippocampus** | | | |
| ***young*** | | | |
| mean [x y z] | **[26.1 -13.6 -17.4]** | **[25.8 -13.6 -17.4]** | **[25.8 -13.3 -17.7]** |
| ± SD | ± [2.18 2.61 1.91] | ± [1.64 2.09 1.40] | ± [1.46 2.04 1.54] |
| size (voxels) ± SD | **17.9** (± 12.23) | **27.0** (± 16.74) | **23.3** (±15.40) |
| ***older*** | | | |
| mean [x y z] | – | **[26.0 -13.1 -17.6]** | **[25.8 -13.2 -17.6]** |
| ± SD | | ± [1.59 2.29 1.49] | [± 1.82 2.02 1.40] |
| size (voxels) ± SD | – | **27.5** (± 14.99) | **23.8** (± 12.27) |
| **Precuneus** | | | |
| ***young*** | | | |
| mean [x y z] | **[7.1 -63.0 38.6]** | **[7.5 -62.6 38.6]** | **[7.3 -62.7 38.6]** |
| ± SD | ± [0.53 1.19 0.92] | ± [0.46 0.92 0.65] | ± [0.36 0.77 0.93] |
| size (voxels) ± SD | **159.4** (±38.53) | **183.7** (± 32.39) | **179.6** (± 25.54) |
| ***older*** | | | |
| mean [x y z] | – | **[7.4 -62.8 38.6]** | **[7.4 -62.7 38.6]** |
| ± SD | | ± [0.37 1.18 1.10] | ± [0.53 1.15 1.11] |
| size (voxels) ± SD | – | **169.1** (± 39.55) | **159.4** (± 39.75) |

Mean ROI coordinates and sizes are shown separately for each cohort and further separated by age group in cohorts 2 and 3. Group averages of the mean x y and z coordinates are shown. PPA: parahippocampal place area.

**Table S9.** *Weighted correlation coefficients and Bayes factors of brain-behavior correlations in older adults.*
(related to Figure 5)

| | A' | VLMT_1to5 | VLMT_Dist | VLMT_5min | VLMT_30min | VLMT_1d | WMS_learn | WMS_30min | WMS_1d |
|---|---|---|---|---|---|---|---|---|---|
| **A-matrix** | | | | | | | | | |
| **PPA-PPA** | $\Pi$ = 0.06<br>$BF_{10}$ = 0.09 | $\Pi$ = 0.25<br>$BF_{10}$ = 8.21 | $\Pi$ = 0.13<br>$BF_{10}$ = 0.24 | $\Pi$ = 0.15<br>$BF_{10}$ = 0.39 | $\Pi$ = 0.16<br>$BF_{10}$ = 0.48 | $\Pi$ = 0.14<br>$BF_{10}$ = 0.30 | $\Pi$ = 0.23<br>$BF_{10}$ = 4.69 | $\Pi$ = 0.14<br>$BF_{10}$ = 0.30 | $\Pi$ = 0.18<br>$BF_{10}$ = 0.86 |
| **PPA-HC** | $\Pi$ = 0.12<br>$BF_{10}$ = 0.21 | $\Pi$ = 0.08<br>$BF_{10}$ = 0.10 | $\Pi$ = -0.17<br>$BF_{10}$ = 0.59 | $\Pi$ = 0.06<br>$BF_{10}$ = 0.08 | $\Pi$ = 0.05<br>$BF_{10}$ = 0.08 | $\Pi$ = 0.10<br>$BF_{10}$ = 0.13 | $\Pi$ = -0.06<br>$BF_{10}$ = 0.09 | $\Pi$ = -0.06<br>$BF_{10}$ = 0.08 | $\Pi$ = -0.02<br>$BF_{10}$ = 0.07 |
| **PPA-Prc** | **$\Pi$ = -0.33<br>$BF_{10}$ = 400.72** | $\Pi$ = -0.08<br>$BF_{10}$ = 0.11 | $\Pi$ = 0.00<br>$BF_{10}$ = 0.07 | $\Pi$ = -0.02<br>$BF_{10}$ = 0.07 | $\Pi$ = -0.03<br>$BF_{10}$ = 0.07 | $\Pi$ = -0.08<br>$BF_{10}$ = 0.10 | $\Pi$ = -0.16<br>$BF_{10}$ = 0.48 | $\Pi$ = -0.10<br>$BF_{10}$ = 0.14 | $\Pi$ = -0.10<br>$BF_{10}$ = 0.13 |
| **HC-PPA** | $\Pi$ = -0.06<br>$BF_{10}$ = 0.08 | $\Pi$ = 0.05<br>$BF_{10}$ = 0.08 | $\Pi$ = -0.01<br>$BF_{10}$ = 0.06 | $\Pi$ = 0.06<br>$BF_{10}$ = 0.09 | $\Pi$ = 0.11<br>$BF_{10}$ = 0.16 | $\Pi$ = 0.11<br>$BF_{10}$ = 0.16 | $\Pi$ = -0.11<br>$BF_{10}$ = 0.16 | $\Pi$ = -0.09<br>$BF_{10}$ = 0.13 | $\Pi$ = -0.15<br>$BF_{10}$ = 0.36 |
| **HC-HC** | $\Pi$ = 0.10<br>$BF_{10}$ = 0.14 | $\Pi$ = -0.00<br>$BF_{10}$ = 0.06 | $\Pi$ = 0.02<br>$BF_{10}$ = 0.07 | $\Pi$ = -0.10<br>$BF_{10}$ = 0.15 | $\Pi$ = -0.02<br>$BF_{10}$ = 0.07 | $\Pi$ = -0.02<br>$BF_{10}$ = 0.06 | $\Pi$ = 0.07<br>$BF_{10}$ = 0.10 | $\Pi$ = 0.04<br>$BF_{10}$ = 0.07 | $\Pi$ = 0.03<br>$BF_{10}$ = 0.07 |
| **HC-Prc** | $\Pi$ = 0.19<br>$BF_{10}$ = 1.23 | $\Pi$ = -0.07<br>$BF_{10}$ = 0.09 | $\Pi$ = -0.24<br>$BF_{10}$ = 6.16 | $\Pi$ = -0.02<br>$BF_{10}$ = 0.07 | $\Pi$ = -0.04<br>$BF_{10}$ = 0.07 | $\Pi$ = -0.14<br>$BF_{10}$ = 0.28 | $\Pi$ = 0.00<br>$BF_{10}$ = 0.06 | $\Pi$ = -0.06<br>$BF_{10}$ = 0.08 | $\Pi$ = -0.08<br>$BF_{10}$ = 0.11 |
| **Prc-PPA** | $\Pi$ = 0.02<br>$BF_{10}$ = 0.07 | $\Pi$ = -0.01<br>$BF_{10}$ = 0.06 | $\Pi$ = -0.09<br>$BF_{10}$ = 0.11 | $\Pi$ = 0.06<br>$BF_{10}$ = 0.08 | $\Pi$ = -0.00<br>$BF_{10}$ = 0.06 | $\Pi$ = 0.06<br>$BF_{10}$ = 0.08 | $\Pi$ = 0.10<br>$BF_{10}$ = 0.14 | $\Pi$ = 0.13<br>$BF_{10}$ = 0.22 | $\Pi$ = 0.16<br>$BF_{10}$ = 0.43 |
| **Prc-HC** | $\Pi$ = -0.16<br>$BF_{10}$ = 0.49 | $\Pi$ = 0.16<br>$BF_{10}$ = 0.47 | $\Pi$ = 0.13<br>$BF_{10}$ = 0.25 | $\Pi$ = 0.07<br>$BF_{10}$ = 0.09 | $\Pi$ = 0.11<br>$BF_{10}$ = 0.16 | $\Pi$ = 0.06<br>$BF_{10}$ = 0.08 | $\Pi$ = -0.10<br>$BF_{10}$ = 0.14 | $\Pi$ = -0.10<br>$BF_{10}$ = 0.14 | $\Pi$ = -0.09<br>$BF_{10}$ = 0.11 |
| **Prc-Prc** | $\Pi$ = -0.01<br>$BF_{10}$ = 0.06 | $\Pi$ = 0.07<br>$BF_{10}$ = 0.09 | $\Pi$ = 0.12<br>$BF_{10}$ = 0.20 | $\Pi$ = -0.04<br>$BF_{10}$ = 0.07 | $\Pi$ = -0.04<br>$BF_{10}$ = 0.07 | $\Pi$ = -0.00<br>$BF_{10}$ = 0.06 | $\Pi$ = -0.05<br>$BF_{10}$ = 0.08 | $\Pi$ = -0.01<br>$BF_{10}$ = 0.06 | $\Pi$ = -0.01<br>$BF_{10}$ = 0.06 |
| **B-matrix** | | | | | | | | | |
| **MEM_PPA-HC** | $\Pi$ = -0.17<br>$BF_{10}$ = 0.58 | $\Pi$ = -0.04<br>$BF_{10}$ = 0.07 | $\Pi$ = -0.02<br>$BF_{10}$ = 0.07 | $\Pi$ = 0.02<br>$BF_{10}$ = 0.07 | $\Pi$ = -0.02<br>$BF_{10}$ = 0.07 | $\Pi$ = -0.06<br>$BF_{10}$ = 0.08 | $\Pi$ = -0.06<br>$BF_{10}$ = 0.08 | $\Pi$ = -0.12<br>$BF_{10}$ = 0.19 | $\Pi$ = -0.10<br>$BF_{10}$ = 0.13 |
| **MEM_PPA-Prc** | $\Pi$ = -0.16<br>$BF_{10}$ = 0.47 | $\Pi$ = -0.05<br>$BF_{10}$ = 0.08 | $\Pi$ = -0.06<br>$BF_{10}$ = 0.08 | $\Pi$ = -0.05<br>$BF_{10}$ = 0.08 | $\Pi$ = -0.10<br>$BF_{10}$ = 0.13 | $\Pi$ = -0.05<br>$BF_{10}$ = 0.08 | $\Pi$ = -0.02<br>$BF_{10}$ = 0.07 | $\Pi$ = 0.02<br>$BF_{10}$ = 0.07 | $\Pi$ = 0.07<br>$BF_{10}$ = 0.10 |
| **MEM_HC-PPA** | $\Pi$ = -0.15<br>$BF_{10}$ = 0.32 | $\Pi$ = -0.01<br>$BF_{10}$ = 0.06 | $\Pi$ = 0.15<br>$BF_{10}$ = 0.38 | $\Pi$ = -0.03<br>$BF_{10}$ = 0.07 | $\Pi$ = -0.04<br>$BF_{10}$ = 0.07 | $\Pi$ = -0.06<br>$BF_{10}$ = 0.08 | $\Pi$ = -0.03<br>$BF_{10}$ = 0.07 | $\Pi$ = -0.06<br>$BF_{10}$ = 0.09 | $\Pi$ = -0.01<br>$BF_{10}$ = 0.07 |
| **MEM_HC-Prc** | $\Pi$ = 0.03<br>$BF_{10}$ = 0.07 | $\Pi$ = 0.03<br>$BF_{10}$ = 0.07 | $\Pi$ = 0.08<br>$BF_{10}$ = 0.11 | $\Pi$ = 0.03<br>$BF_{10}$ = 0.07 | $\Pi$ = 0.00<br>$BF_{10}$ = 0.06 | $\Pi$ = -0.10<br>$BF_{10}$ = 0.13 | $\Pi$ = 0.08<br>$BF_{10}$ = 0.10 | $\Pi$ = 0.04<br>$BF_{10}$ = 0.07 | $\Pi$ = 0.02<br>$BF_{10}$ = 0.07 |
| **MEM_Prc-PPA** | $\Pi$ = -0.03<br>$BF_{10}$ = 0.07 | $\Pi$ = 0.01<br>$BF_{10}$ = 0.06 | $\Pi$ = 0.09<br>$BF_{10}$ = 0.11 | $\Pi$ = 0.06<br>$BF_{10}$ = 0.08 | $\Pi$ = 0.08<br>$BF_{10}$ = 0.11 | $\Pi$ = 0.11<br>$BF_{10}$ = 0.16 | $\Pi$ = 0.02<br>$BF_{10}$ = 0.07 | $\Pi$ = 0.13<br>$BF_{10}$ = 0.24 | $\Pi$ = 0.10<br>$BF_{10}$ = 0.13 |
| **MEM_Prc-HC** | $\Pi$ = 0.13<br>$BF_{10}$ = 0.22 | $\Pi$ = 0.07<br>$BF_{10}$ = 0.09 | $\Pi$ = -0.01<br>$BF_{10}$ = 0.06 | $\Pi$ = 0.07<br>$BF_{10}$ = 0.10 | $\Pi$ = 0.09<br>$BF_{10}$ = 0.11 | $\Pi$ = 0.18<br>$BF_{10}$ = 0.73 | $\Pi$ = 0.03<br>$BF_{10}$ = 0.07 | $\Pi$ = 0.00<br>$BF_{10}$ = 0.06 | $\Pi$ = 0.06<br>$BF_{10}$ = 0.09 |
| **C-matrix** | | | | | | | | | |
| **Input_PPA** | $\Pi$ = 0.19<br>$BF_{10}$ = 1.19 | **$\Pi$ = 0.26<br>$BF_{10}$ = 17.74** | $\Pi$ = 0.18<br>$BF_{10}$ = 0.68 | $\Pi$ = 0.23<br>$BF_{10}$ = 4.54 | **$\Pi$ = 0.26<br>$BF_{10}$ = 15.65** | $\Pi$ = 0.22<br>$BF_{10}$ = 3.52 | **$\Pi$ = 0.27<br>$BF_{10}$ = 19.79** | **$\Pi$ = 0.26<br>$BF_{10}$ = 12.77** | **$\Pi$ = 0.27<br>$BF_{10}$ = 15.08** |

The table displays Shepherd's *Pi* correlation coefficients averaged across cohorts 2 and 3 and weighted by the effective sample sizes (i.e., after outlier exclusions). Cumulative Bayes factors calculated as the Bayes factors for the weighted mean correlation coefficients, based on the total effective sample size. Correlations with a cumulative Bayes factor > 10 are highlighted.

**Table S10.** *Weighted correlation coefficients and Bayes factors of brain-behavior correlations in young adults (cohorts 1, 2, and 3).*
(related to Figure 5)

| | A‘ | VLMT_1to5 | VLMT_Dist | VLMT_5min | VLMT_30min | VLMT_1d | WMS_learn | WMS_30min | WMS_1d |
|---|---|---|---|---|---|---|---|---|---|
| **A-matrix** | | | | | | | | | |
| **PPA-PPA** | $\Pi$ = -0.02<br>$BF_{10}$ = 0.06 | $\Pi$ = 0.14<br>$BF_{10}$ = 0.36 | $\Pi$ = 0.10<br>$BF_{10}$ = 0.16 | $\Pi$ = -0.08<br>$BF_{10}$ = 0.10 | $\Pi$ = -0.02<br>$BF_{10}$ = 0.06 | $\Pi$ = -0.11<br>$BF_{10}$ = 0.18 | $\Pi$ = -0.03<br>$BF_{10}$ = 0.06 | $\Pi$ = -0.05<br>$BF_{10}$ = 0.07 | $\Pi$ = -0.06<br>$BF_{10}$ = 0.08 |
| **PPA-HC** | $\Pi$ = 0.12<br>$BF_{10}$ = 0.30 | $\Pi$ = 0.09<br>$BF_{10}$ = 0.12 | $\Pi$ = 0.05<br>$BF_{10}$ = 0.07 | $\Pi$ = 0.01<br>$BF_{10}$ = 0.06 | $\Pi$ = 0.11<br>$BF_{10}$ = 0.17 | $\Pi$ = 0.07<br>$BF_{10}$ = 0.10 | $\Pi$ = 0.12<br>$BF_{10}$ = 0.24 | $\Pi$ = 0.11<br>$BF_{10}$ = 0.19 | $\Pi$ = 0.16<br>$BF_{10}$ = 0.74 |
| **PPA-Prc** | $\Pi$ = -0.20<br>$BF_{10}$ = 3.96 | $\Pi$ = -0.15<br>$BF_{10}$ = 0.59 | $\Pi$ = -0.17<br>$BF_{10}$ = 1.28 | $\Pi$ = -0.09<br>$BF_{10}$ = 0.13 | $\Pi$ = -0.14<br>$BF_{10}$ = 0.41 | $\Pi$ = -0.15<br>$BF_{10}$ = 0.50 | $\Pi$ = -0.21<br>$BF_{10}$ = 4.76 | $\Pi$ = -0.20<br>$BF_{10}$ = 3.20 | $\Pi$ = -0.17<br>$BF_{10}$ = 1.18 |
| **HC-PPA** | $\Pi$ = -0.06<br>$BF_{10}$ = 0.07 | $\Pi$ = -0.12<br>$BF_{10}$ = 0.24 | $\Pi$ = -0.11<br>$BF_{10}$ = 0.18 | $\Pi$ = -0.13<br>$BF_{10}$ = 0.29 | $\Pi$ = -0.03<br>$BF_{10}$ = 0.06 | $\Pi$ = -0.08<br>$BF_{10}$ = 0.10 | $\Pi$ = -0.10<br>$BF_{10}$ = 0.17 | $\Pi$ = -0.11<br>$BF_{10}$ = 0.19 | $\Pi$ = -0.14<br>$BF_{10}$ = 0.45 |
| **HC-HC** | $\Pi$ = -0.16<br>$BF_{10}$ = 0.97 | $\Pi$ = 0.03<br>$BF_{10}$ = 0.06 | $\Pi$ = -0.04<br>$BF_{10}$ = 0.06 | $\Pi$ = 0.09<br>$BF_{10}$ = 0.13 | $\Pi$ = 0.11<br>$BF_{10}$ = 0.18 | $\Pi$ = -0.01<br>$BF_{10}$ = 0.06 | $\Pi$ = -0.02<br>$BF_{10}$ = 0.06 | $\Pi$ = -0.10<br>$BF_{10}$ = 0.15 | $\Pi$ = -0.09<br>$BF_{10}$ = 0.13 |
| **HC-Prc** | $\Pi$ = 0.05<br>$BF_{10}$ = 0.07 | $\Pi$ = 0.08<br>$BF_{10}$ = 0.11 | $\Pi$ = -0.05<br>$BF_{10}$ = 0.07 | $\Pi$ = 0.06<br>$BF_{10}$ = 0.08 | $\Pi$ = -0.06<br>$BF_{10}$ = 0.08 | $\Pi$ = -0.03<br>$BF_{10}$ = 0.06 | $\Pi$ = 0.16<br>$BF_{10}$ = 0.85 | $\Pi$ = 0.19<br>$BF_{10}$ = 2.28 | $\Pi$ = 0.21<br>$BF_{10}$ = 4.58 |
| **Prc-PPA** | $\Pi$ = -0.08<br>$BF_{10}$ = 0.11 | $\Pi$ = -0.05<br>$BF_{10}$ = 0.07 | $\Pi$ = -0.00<br>$BF_{10}$ = 0.05 | $\Pi$ = 0.14<br>$BF_{10}$ = 0.40 | $\Pi$ = 0.12<br>$BF_{10}$ = 0.25 | $\Pi$ = 0.15<br>$BF_{10}$ = 0.55 | $\Pi$ = 0.04<br>$BF_{10}$ = 0.07 | $\Pi$ = 0.07<br>$BF_{10}$ = 0.09 | $\Pi$ = 0.03<br>$BF_{10}$ = 0.06 |
| **Prc-HC** | $\Pi$ = -0.07<br>$BF_{10}$ = 0.10 | $\Pi$ = -0.12<br>$BF_{10}$ = 0.26 | $\Pi$ = -0.16<br>$BF_{10}$ = 0.83 | $\Pi$ = -0.01<br>$BF_{10}$ = 0.06 | $\Pi$ = -0.00<br>$BF_{10}$ = 0.06 | $\Pi$ = 0.04<br>$BF_{10}$ = 0.06 | $\Pi$ = 0.06<br>$BF_{10}$ = 0.08 | $\Pi$ = 0.08<br>$BF_{10}$ = 0.11 | $\Pi$ = 0.11<br>$BF_{10}$ = 0.20 |
| **Prc-Prc** | $\Pi$ = 0.15<br>$BF_{10}$ = 0.58 | $\Pi$ = 0.02<br>$BF_{10}$ = 0.06 | $\Pi$ = 0.08<br>$BF_{10}$ = 0.11 | $\Pi$ = 0.07<br>$BF_{10}$ = 0.09 | $\Pi$ = 0.03<br>$BF_{10}$ = 0.06 | $\Pi$ = 0.01<br>$BF_{10}$ = 0.06 | $\Pi$ = -0.05<br>$BF_{10}$ = 0.07 | $\Pi$ = -0.06<br>$BF_{10}$ = 0.08 | $\Pi$ = -0.07<br>$BF_{10}$ = 0.09 |
| **B-matrix** | | | | | | | | | |
| **MEM_PPA-HC** | $\Pi$ = 0.10<br>$BF_{10}$ = 0.17 | $\Pi$ = 0.03<br>$BF_{10}$ = 0.06 | $\Pi$ = 0.01<br>$BF_{10}$ = 0.06 | $\Pi$ = -0.04<br>$BF_{10}$ = 0.06 | $\Pi$ = 0.02<br>$BF_{10}$ = 0.06 | $\Pi$ = -0.04<br>$BF_{10}$ = 0.07 | $\Pi$ = -0.03<br>$BF_{10}$ = 0.06 | $\Pi$ = -0.04<br>$BF_{10}$ = 0.07 | $\Pi$ = -0.07<br>$BF_{10}$ = 0.09 |
| **MEM_PPA-Prc** | $\Pi$ = 0.01<br>$BF_{10}$ = 0.05 | $\Pi$ = 0.09<br>$BF_{10}$ = 0.13 | $\Pi$ = 0.04<br>$BF_{10}$ = 0.07 | $\Pi$ = 0.22<br>$BF_{10}$ = 8.56 | $\Pi$ = 0.17<br>$BF_{10}$ = 0.96 | **$\Pi$ = 0.25**<br>**$BF_{10}$ = 35.87** | $\Pi$ = 0.18<br>$BF_{10}$ = 1.49 | $\Pi$ = 0.18<br>$BF_{10}$ = 1.48 | $\Pi$ = 0.14<br>$BF_{10}$ = 0.41 |
| **MEM_HC-PPA** | $\Pi$ = -0.03<br>$BF_{10}$ = 10.06 | $\Pi$ = 0.02<br>$BF_{10}$ = 0.06 | $\Pi$ = -0.05<br>$BF_{10}$ = 0.07 | $\Pi$ = 0.00<br>$BF_{10}$ = 0.06 | $\Pi$ = 0.01<br>$BF_{10}$ = 0.06 | $\Pi$ = -0.04<br>$BF_{10}$ = 0.07 | $\Pi$ = -0.07<br>$BF_{10}$ = 0.09 | $\Pi$ = -0.06<br>$BF_{10}$ = 0.08 | $\Pi$ = -0.04<br>$BF_{10}$ = 0.07 |
| **MEM_HC-Prc** | $\Pi$ = -0.04<br>$BF_{10}$ = 0.06 | $\Pi$ = 0.02<br>$BF_{10}$ = 0.06 | $\Pi$ = -0.02<br>$BF_{10}$ = 0.06 | $\Pi$ = -0.04<br>$BF_{10}$ = 0.06 | $\Pi$ = -0.05<br>$BF_{10}$ = 0.07 | $\Pi$ = -0.07<br>$BF_{10}$ = 0.09 | $\Pi$ = -0.01<br>$BF_{10}$ = 0.06 | $\Pi$ = 0.04<br>$BF_{10}$ = 0.06 | $\Pi$ = 0.03<br>$BF_{10}$ = 0.06 |
| **MEM_Prc-PPA** | $\Pi$ = -0.03<br>$BF_{10}$ = 0.06 | $\Pi$ = 0.12<br>$BF_{10}$ = 0.26 | $\Pi$ = -0.00<br>$BF_{10}$ = 0.05 | $\Pi$ = 0.10<br>$BF_{10}$ = 0.14 | $\Pi$ = 0.09<br>$BF_{10}$ = 0.13 | $\Pi$ = 0.06<br>$BF_{10}$ = 0.08 | $\Pi$ = -0.05<br>$BF_{10}$ = 0.07 | $\Pi$ = -0.04<br>$BF_{10}$ = 0.07 | $\Pi$ = -0.07<br>$BF_{10}$ = 0.09 |
| **MEM_Prc-HC** | $\Pi$ = -0.01<br>$BF_{10}$ = 0.05 | $\Pi$ = 0.13<br>$BF_{10}$ = 0.34 | $\Pi$ = -0.04<br>$BF_{10}$ = 0.06 | $\Pi$ = 0.03<br>$BF_{10}$ = 0.06 | $\Pi$ = 0.05<br>$BF_{10}$ = 0.07 | $\Pi$ = 0.09<br>$BF_{10}$ = 0.13 | $\Pi$ = -0.07<br>$BF_{10}$ = 0.09 | $\Pi$ = -0.05<br>$BF_{10}$ = 0.07 | $\Pi$ = -0.10<br>$BF_{10}$ = 0.15 |
| **C-matrix** | | | | | | | | | |
| **Input_PPA** | **$\Pi$ = 0.31**<br>**$BF_{10}$ = 5248.48** | $\Pi$ = 0.14<br>$BF_{10}$ = 0.47 | $\Pi$ = 0.02<br>$BF_{10}$ = 0.06 | $\Pi$ = 0.05<br>$BF_{10}$ = 0.07 | $\Pi$ = 0.03<br>$BF_{10}$ = 0.06 | $\Pi$ = 0.03<br>$BF_{10}$ = 0.06 | $\Pi$ = 0.04<br>$BF_{10}$ = 0.06 | $\Pi$ = 0.10<br>$BF_{10}$ = 0.16 | $\Pi$ = 0.06<br>$BF_{10}$ = 0.08 |

The table displays Shepherd's *Pi* correlation coefficients averaged across cohorts 1, 2 and 3 and weighted by the effective sample sizes (i.e., after outlier exclusions). Cumulative Bayes factors calculated as the Bayes factors for the weighted mean correlation coefficients, based on the total effective sample size. Correlations with a cumulative Bayes factor > 10 are highlighted.
